# Mathematical Modeling and Analysis of Mitochondrial Retrograde Signaling Dynamics

**DOI:** 10.1101/2021.03.27.437239

**Authors:** Shao-Ting Chiu, Wen-Wei Tseng, An-Chi Wei

## Abstract

Mitochondria, semi-autonomous eukaryotic organelles, participate in energy production and metabolism, making mitochondrial quality control crucial. As most mitochondrial proteins are encoded by nuclear genes, quality control depends on proper mitochondria-nucleus communication, designated mitochondrial retrograde signaling. Early studies focused on retrograde signaling participants and specific gene knockouts. However, mitochondrial signal modulation remains elusive. Using yeast, we simulated signal propagation following mitochondrial damage and proposed a mathematical model based ordinary differential equations. Mitochondrial retrograde signaling decisions were described using a Boolean model. Dynamics were analyzed through an ordinary differential equation-based model and extended to evaluate the model response to noisy damage signals. Simulation revealed localized protein concentration dynamics, including waveforms, frequency response, and robustness under noise. Retrograde signaling is bistable with three localized steady states, and increased damage compromises robustness. We elucidated mitochondrial retrograde signaling, thus providing a basis for drug design against yeast and fungi.

**Author Summary:** The yeast RTG pathway regulates mitochondrial metabolism and mitochondrial quality through passing mitochondrial signal to the nucleus to modulate gene expressions. Using microscopic data of RTG proteins translocation from cytosol to nucleus, the parameters were found by fitting 16 knockout conditions simulated from the proposed differential equation-based model. Further, through dose response, ultrasensitivity, frequency response and noise interference, we demonstrated a switch-like property of RTG pathway activation, a capacity charging property of RTG protein translocation, a low pass filter property of signals in different frequencies, and noise amplification due to mitochondrial signal. The proposed mathematical model enables us to understand the dynamics and mechanisms in mitochondrial retrograde signaling in yeast and provides potential antifungal treatment strategies.

## Introduction

Mitochondria serve as the cells’ powerhouse, utilizing the established proton motive force to generate the high-energy molecule, ATP, thus facilitating various cellular processes [1]. The analogy of a simple electrical circuit model is a convenient way to describe the interactions between ion dynamics and energetics in the complex mitochondrial system [2, 3]. Mitochondria regulate various cellular mechanisms under normal physiological conditions, as well as in disease, including ATP generation, regulation of signal transduction, immune responses, and cell apoptosis [1]. Therefore, mitochondria serve as an essential regulatory metabolic hub [4].

However, their extensive role within the cell renders mitochondria prone to be damaged [5, 6]. Mitochondrial quality control is therefore essential for cell viability, which relies on mitochondria-driven energy production [1], lipid synthesis [7], and amino acid metabolism [8]. However, energy production via the electron transport chain results in an oxidative stress burden [9]. Unlike other membrane-bound organelles, which rely solely on protein products from the nuclear genome, mitochondria possess their own genome, encoding dedicated enzymes and repair systems; however, they remain semi-autonomous organelles, as most mitochondria protein-encoding genes are translocated into the nucleus [10]. In fact, in budding yeast, only 35 of the 1,000 mitochondrial genes are located in the mitochondrial genome, representing a minor fraction of the metabolic system [11]. Therefore, mitochondria require various products from nucleus genome to maintain functionality. The interaction between mitochondria and the nucleus is a bidirectional process [4, 12]. Once mitochondria are damaged, their status is reported to the nucleus via signaling factors or metabolic pathways, enabling the cell to coordinate mitochondrial biogenesis. Furthermore, mitochondrial quality control depends on mitochondrial feedback, referred to as mitochondrial retrograde signaling [5, 12–14].

The most well-characterized mitochondrial RTG signaling cascade is that in budding yeast [12, 15–18], which gets triggered when mitochondrial respiratory function is compromised [19], causing a shift in carbohydrate and nitrogen metabolism [20]. *CIT2*, one of the regulated genes, encodes a peroxisomal isoform of citrate syntheses involved in the glyoxylate cycle, thereby bypassing the TCA cycle. Thus, the upregulation of *CIT2* expression allows for more efficient usage of carbon sources when mitochondria are damaged. This RTG pathway was first characterized by investigating various yeast gene deletion strains. For instance, in a study by Sekito et al. [18], RTG genes were independently knocked out, and green fluorescent protein (GFP) was employed to localize the RTG proteins, revealing details of the mitochondria-to-nucleus communication system. Furthermore, the localization of RTG proteins was compared among deletion strains, as well as strains with dysfunctional mitochondria, such as *ρ*^0^ cells, which lack a mitochondrial genome.

The RTG pathway is initiated by the activation of Rtg2p, a cytoplasmic sensor of mitochondrial damage with an ATP-binding domain similar to the Hsp70/actin/sugar kinase superfamily. Rtg2p is activated by the loss of mitochondrial membrane potential or the ATP/ADP ratio [12, 20, 21], which results in the degradation of Bmhp/Mksp heterodimer that inhibits the Rtg3p activation [17, 21]. Rtg1p/3p are are basic helix-loop-helix-leucine zipper (bHLH/Zip) transcription factors that induce the retrograde response via nucleus translocation with nucleus localization signal (NLS) and binding to the promoter R box region of RTG response genes to initiate transcription [22].

Without Rtg2p sensing mitochondrial status, Rtg3p and Rtg1p localization occurs independently of mitochondrial damage because of constant inhibition of Mksp. Rtg3p and Rtg1p remain cytoplasmic in wild type and dysfunctional mitochondria (*ρ*^0^ strain) [18]. In addition, Rtg1p regulates the translocation of Rtg3p. Although Rtg1p participates in the activation of the retrograde response, it retains Rtg3p in the cytoplasm in the absence of mitochondrial damage. In contrast, Rtg3p accumulates in the nucleus regardless of mitochondrial status in Rtg1p knockout mutants, while the opposite is not true. Initiation of retrograde response signaling requires nuclear accumulation of both Rtg3p and Rtg1p like the AND logic gate [18, 23, 24]. Therefore, Rtg1p acts as a positive regulator of retrograde response genes, as well as a negative regulator that retains Rtg3p in the cytoplasm in the absence of mitochondrial damage.

Cells usually harbor a number of mitochondria that share the same retrograde signaling pathway, resulting in a multiplexing issue first described in telecommunications [25]. Sharing signaling channels may lead to unwanted crosstalk. Further, biochemical pathway noise restricts the cells’ ability to receive information from their organelles. In eukaryotic cells, such as budding yeast, mitochondria interact with the nucleus via the same biochemical channels, and the consequent multiplexing issue represents a barrier to assessing the health state of each mitochondrion. How biological systems manage multiplex communication networks during the mitochondrial RTG response remains unclear.

To investigate the communication properties of mitochondrial retrograde signaling, we have developed a differential equation-based model to simulate the process. The mitochondrial status and deleted genes are defined as input, whereas translocation of Rtg1p/Rtg3p is the defined output. These interactions were previously observed using immunofluorescence microscopy [18, 32], and subsequently summarized in a Boolean table. The parameters were fitted with the Boolean relationship using Monte Carlo simulation. A parameter set was verified by solving the steady states under each observed genotype. Collectively, our mathematical model interprets the dynamics of mitochondrial retrograde signaling. Furthermore, the proposed mathematical model also has potential applications in research on antifungal drugs and aging.

## Results

### A differential equation-based model of mitochondrial retrograde signaling

As a channel of communication between mitochondria and the nucleus, RTG signaling is well regulated and fine-tuned for a proper response to mitochondrial damage signals. The dynamics and interactions of Rtg1p, Rtg2p, Rtg3p, Msk1p, and Bmhp can be described using ordinary differential equations (ODE) and biochemical reaction network. The first part of RTG signaling is the activation layer, during which the mitochondrial damage signal activates Rtg2p. Then, the signal is transmitted to the modulation layer where Rtg1/3p translocate into the nucleus and initiate transcription (Fig 1a). For simplicity, the chemical interactions among retrograde proteins and the translocation of Rtg1/3p are assumed to obey the law of mass action, and the total concentration of each protein was regarded as conserved. To connect retrograde proteins with mitochondrial damage, the activation of Rtg2p was modeled using the Hill equation with the sum of mitochondrial damage. Furthermore, proteins with activation states or locations are defined separately, leading to a total of 17 protein variables and 24 kinetic coefficients (Table S3).

**Fig 1.**
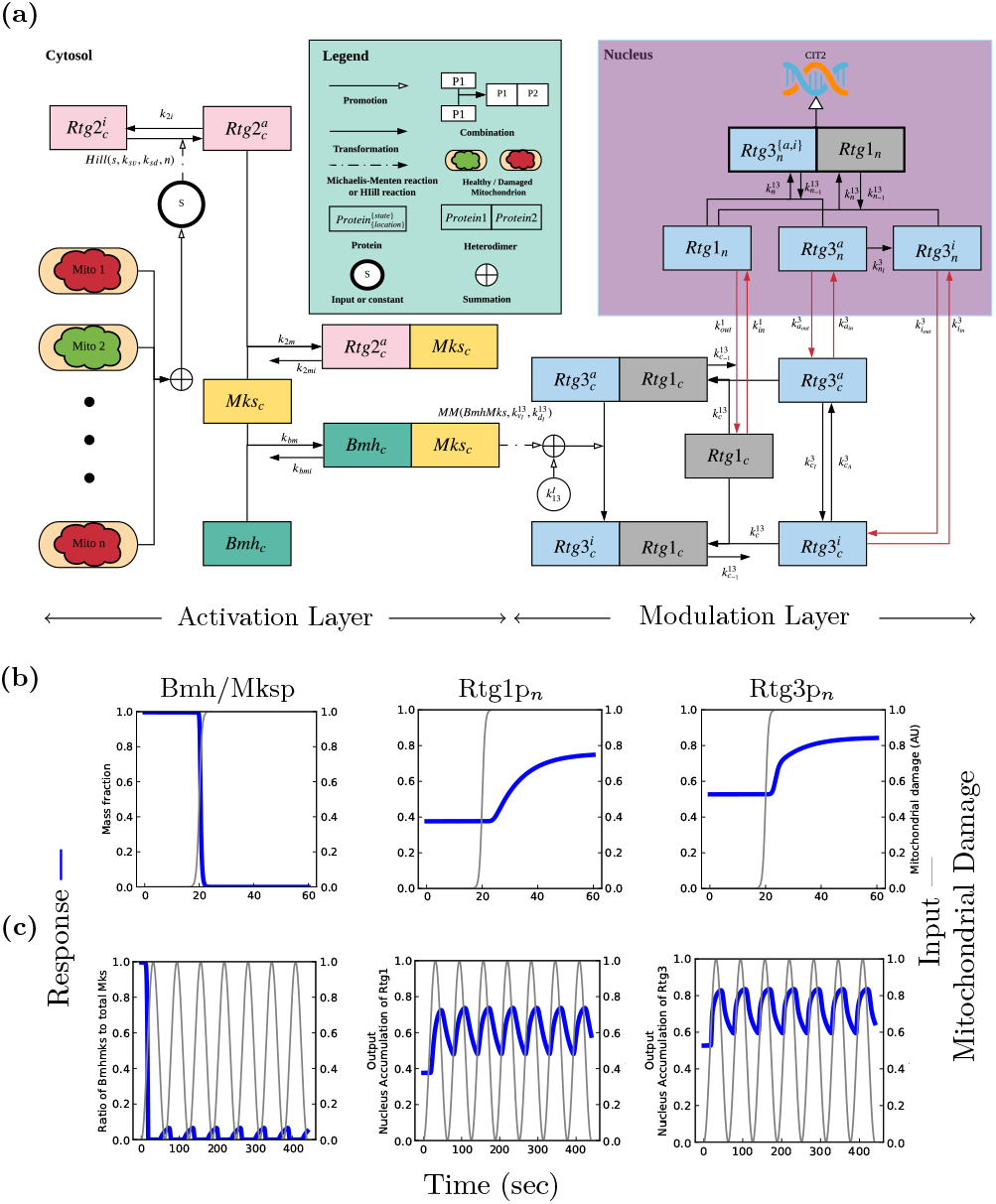
Mitochondrial retrograde signaling pathway model. (a) The mitochondria-to-nucleus communication circuit. Retrograde signaling is divided into the activation layer and modulation layer. Retrograde proteins (Bmhp, Mksp, and Rtg1/2/3p) are classified into an active (*-act*) and inactive state (*-ina*). Cellular compartments are divided based on the nucleus (*-nuc*) and cytosol (*-cyt*). The arrow represents the binding of two proteins, transformation of active states, or translocation between nucleus and cytosol, with the kinetic coefficient labeled besides it. The input s is defined as the summation of mitochondrial damage signals. Rtg2p is activated once the mitochondrion loses its membrane potential, or the ATP concentration decreases. The active form of Rtg2p suppresses the formation of the Mks/bmhp complex by competitive binding with Mksp, resulting in dephosphorylation of Rtg3p and translocates to the nucleus accompanied by Rtg1p. The Rtg1p-Rtg3p complex eventually triggers the retrograde response and upregulates *CIT2* expression. (b) Step responses of Bmh/Mksp, Rtg1p_*n*_ and Rtg3p_*n*_ showing switch-like responses of Bmh/Mksp, capacity-charging curve with a time delay for Rtg1p_*n*_ and delayed two-staged rise with a sharp transition at the beginning, for Rtg3p_*n*_. (c) The sinusoidal inputs of 1.6 × 10^−2^ Hz response reveals RTG signaling pathway as a low-pass filter. The mitochondrial damage signal (thin gray line), labeled as the input with quantity shown on the right y-axis.The concentration of RTG proteins (thick blue line), regarded as outputs with quantities labeled on the left y-axis. To normalize the response, the mass fraction is defined as the ratio of the concentration of a given protein set to its total concentration. Nucleus accumulation is described by the ratio of nucleus concentration to total protein concentration.

To identify the parameter set that would make the model compatible with the experimental observations [18, 32, 34], a Monte Carlo simulation method was used to perform the parameter exploration. First, to leverage the microscopic data [18, 32], the translocation of Rtg1/3p and mitochondrial states were summarized in a Boolean model with 17 genetic backgrounds, including the combination of RTG gene knockouts (Table 1). Second, the relative protein concentrations of Bmhp and Rtg1/2/3p were determined based on the gene expression [35] and quantitative western blot analyses [34](Fig S1). Third, a random search method was applied to explore the parameter set. Finally, the steady states of the model were compared to the Boolean model (Fig 2b). A parameter set was regarded as valid when the steady states fulfilled the observations of 17 knockout conditions.

**Fig 2.**
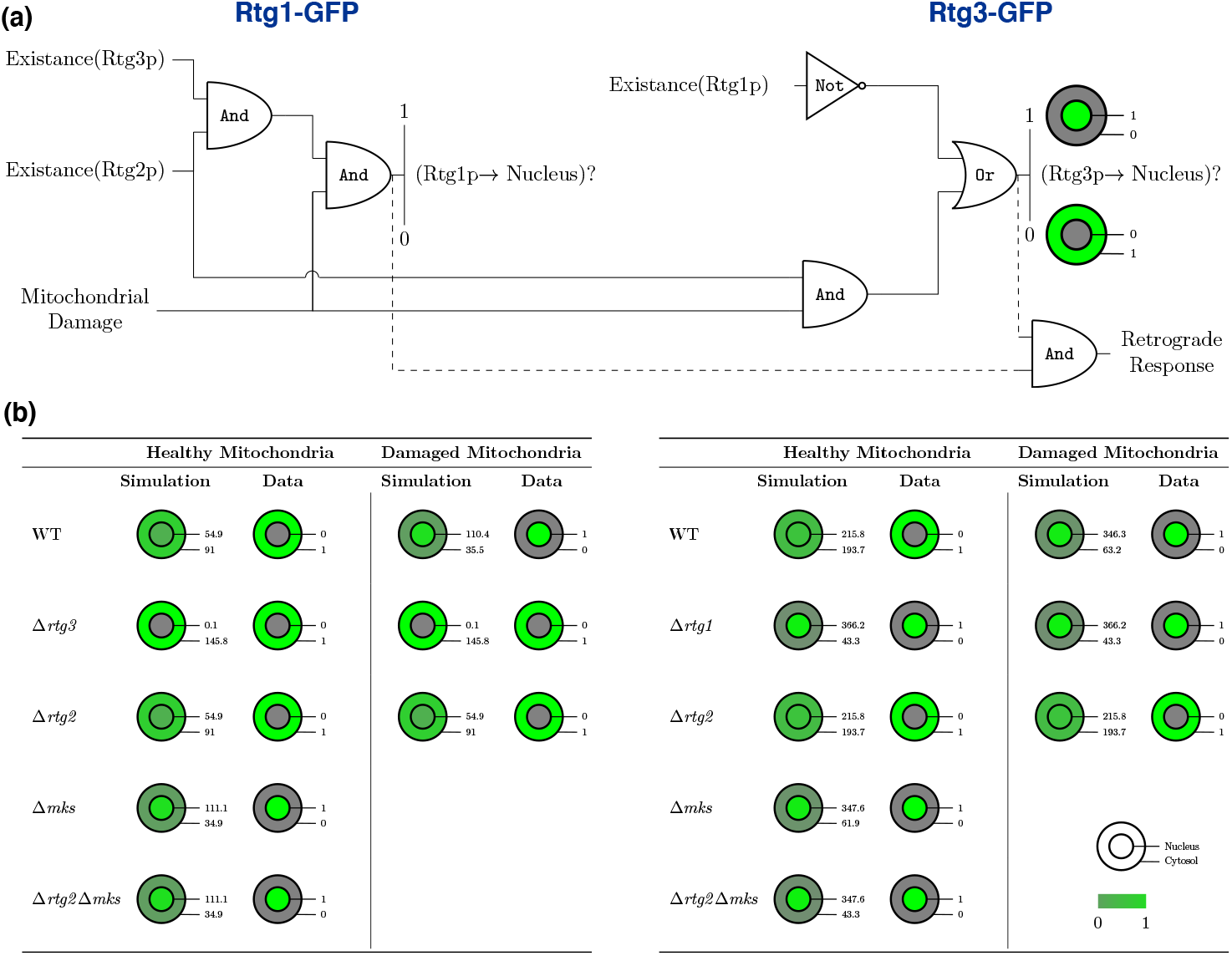
Boolean decision in RTG signaling pathway. (a) Logic gate analog for the RTG signaling pathway Rtg1p(left) and Rtg3p(right). Mitochondrial damage and the existence of Rtg1p, Rtg2p, and Rtg3p determined the translocation of Rtg1p into nucleus through a series of logic gate decisions (Table S2). The input is defined as 0 with knockout protein or absence of mitochondrial damage, and 1 represents the existence of a protein or mitochondrial damage. On the other hand, the output is defined as 1 with nucleus accumulation and 0 means no accumulation event. The solid line represents logic gate decision for nucleus accumulation, and the dotted line represents biological process that leads to retrograde response. (b) Boolean transformation of Rtg proteins nucleus translocation in healthy and damaged mitochondria. Simulated distribution of GFP-labeled Rtg1p under multiple conditions was compared to the Boolean model. The steady states of GFP proteins are listed by the cell icon labeled with nucleus (inner circle) and cytoplasmic (outer circle) concentration in arbitrary units. Localized concentrations are described as the summation of GFP-labeled proteins, including heterodimers. The relative quantity indicates the localized concentration-to-summation ratio, from 0 (gray) to 1 (green). To verify the result, simulated distributions are compared with the qualitative data of Δ*rtg1*/Δ*rtg2*/Δ*rtg3* [18] and Δ*mks* [21]. Nuclear accumulation is regarded as nuclear-to-cytoplasmic ratio is greater than 1.5. Simulation distribution of GFP-labeled Rtg1p(left) and GFP-labeled Rtg3p(right).

**Table 1.**
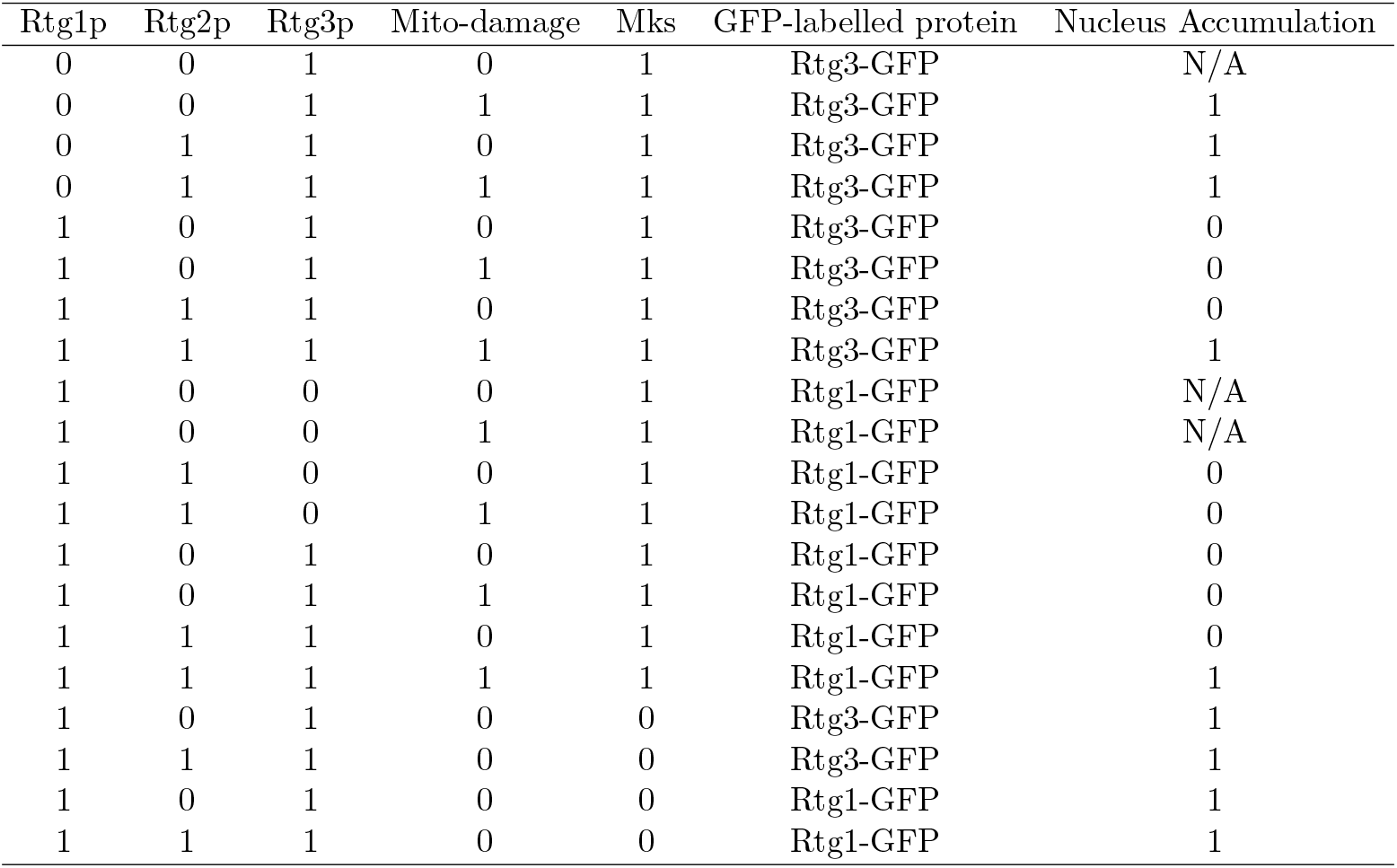
The Boolean model of mitochondrial retrograde signaling. The existence of RTG proteins and mitochondrial damage signal is described in binary quantity based on the experimental observations of the sub-cellular localization of GFP-tagged-RTG proteins in wild type and rtgΔ mutant derivatives of *ρ*_+_ and *ρ*_0_ cells [18]. The 0 represents deleted protein or absence of mitochondrial damage, and 1 represents the opposite phenomena. Untested conditions are designated as N/A.

### The ultrasensitivity of Bmh/Mksp degradation causes a switch-like response of Rtg1/3p translocation to mitochondrial damage

Nuclear accumulation of Rtg1p and Rtg3p activates the retrograde response. To understand the mechanism by which information is propagated from the upstream Bmh/Mksp inhibitor to Rtg1/3p translocation, a sigmoid signal of mitochondrial damage was applied to simulate the transition of RTG proteins. The step response revealed sophisticated details on RTG protein dynamics induced by the fast sigmoid transition of the damage signal (Fig 1b). The degradation of Bmh/Mksp indicated an ultrasensitive response to the input signal. However, the translocation of Rtg1p and Rtg3p was relatively smoother and delayed. The delayed stimulation of Rtg1p and Rtg3p may be caused by the slow Rtg3p autoactivation rate. Besides, the step responses overshoot pattern was identified in the particular forms of Rtg1p and Rtg3p, while summation canceled this effect (Fig S2a).

Two forces control the nuclear translocation of Rtg3p, namely autoactivation of the NLS and degradation of the cytoplasmic Rtg1/3p heterodimer, releasing extra Rtg3p monomers that permeate the nuclear membrane. When the mitochondrial damage signal degrades Bmh/Mksp via Rtg2p, the autoactivation of Rtg3p turns on the NLS and increases the influx kinetic coefficient. Furthermore, the overshoot of activated Rtg3p concentration causes the decline observed in the early response (Fig S2a). The decreased cytoplasmic Rtg3p concentration causes further degradation of the Rtg1/3p heterodimer, releasing more free-Rtg3p. Autoactivation of the Rtg3p NLS contributes to the sharp increase in Rtg3p nuclear concentration shortly after the damage signal. Once autoactivation reaches a steady-state, free Rtg3p is conserved and translocated to the nucleus by simple diffusion. Hence, the transition of Rtg3p becomes smoother in the second stage, leading to a capacity-charging curve at the end (Fig S2b). In addition to Rtg3p, the translocation of Rtg1p also results from degradation of the cytoplasmic Rtg1/3p heterodimer and simple diffusion. When Rtg1p is absent or negligible, simple diffusion causes Rtg3p to accumulate in the nucleus independently of the damage signal. The role of Rtg1p is to retain Rtg3p in the cytoplasm, however, the anchoring force is insufficient to retain activated Rtg3p [18]. The current simulation supports previous observations and provides details regarding the ultrasensitivity of Bmh/Mksp degradation, the two-staged transition of Rtg3p, and the capacity-charging curve of Rtg1p in response to mitochondrial damage signals.

### Competitive binding to Mksp between Bmhp and Rtg2p contributes to the ultrasensitivity of Bmh/Mksp degradation

The ultrasensitivity of Bmh/Mksp degradation is indicated by the phase plot with mitochondrial damage as input (Fig 3). Although the Hill function models this process with the dissociation constant of input(*k_sd_*), the phase plot of Rtg2p reveals a two-staged sigmoid transition with two thresholds significantly below *k_sd_*, and with higher stiffness on the first one. The degradation of Bmh/Mksp occurs at the first transition, wherein the threshold for sensing mitochondrial damage is lower than that of the input Hill function. Therefore, the ultrasensitivity of Bmh/Mksp is independent of the Hill input model (Fig S3) and is caused by the competitive binding between activated Rtg2p and Bmhp.

**Fig 3.**
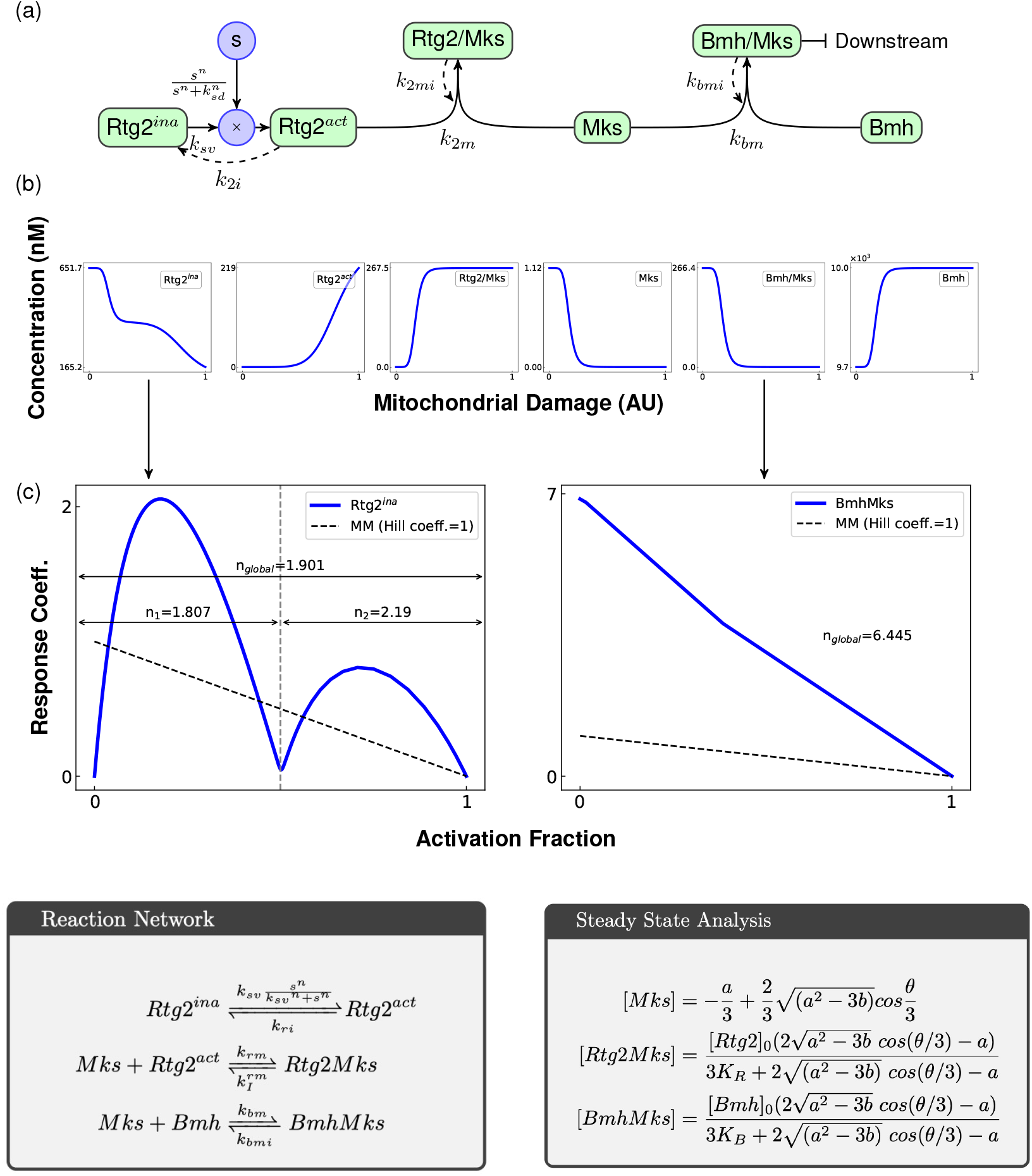
Ultrasensitivity of the Bmh/Mksp degradation results from molecular titration. (a) Scheme of the competitive binding between Rtg2p and Bmhp with Mksp. The forward reactions are labeled with thick arrows, and reverse reactions are labeled with dashed arrows. (b) Phase plots. The input signal is defined within the interval between 0 and 1, and the corresponding steady states are plotted separately with the parameters: *n* = 7, *k_sd_* = 0.96, *k_sv_* = 11.67, *k*_2*i*_ = 4.94, *k*_2*mi*_ = 0.043, *k*_2*m*_ = 1604.15, *k_bmi_* = 2.413, *k_bm_* = 0.059in arbitrary unit.(c) Relative amplification approach of Rtg2p^*ina*^ and Bmh/Mksp in response to mitochondrial damage. The activated fraction is converted from the input signal *s* normalized by its range [36]; the response coefficient is the normalized sensitivity of output with respect to the input signal [57]. The transformed responses of Rtg2p^*ina*^ and Bmh/Mksp (thick blue line) are compared via the Michaelis-Menten reaction with a Hill coefficient equal to one, described as a linear function (dashed line) [36]. *n_global_* represents global Hill coefficient; *n*_1_ and *n*_2_ are local Hill coefficients in the first and second transition of Rtg2p^*ina*^.

To quantify the sensitivity, the relative amplification method [36], a quantitative approach for estimating the Hill coefficient with basal activation, was applied. As shown in Fig 3c, the response of inactive Rtg2p (Rtg2p^*ina*^) is with lower Hill coefficients for both transitions compared to the input locally and globally (Fig 3c, left). Meanwhile, the degradation of the Bmh/Mksp heterodimer occurs at the first transition of Rtg2p^*ina*^, with a higher Hill coefficient, which was lower than that of the input (Fig 3c, right). Via the relative amplification method, we estimated the Hill coefficient numerically. Despite knowing the quantity of the Hill coefficient, the transition of Bmh/Mksp occurred far below the dissociation constant *k_sd_*, which is unlikely to be caused by the input Hill function, leading to questions regarding the source of its ultrasensitivity. To elucidate the underlying mechanism, we analyzed the Rtg2p-Mksp-Bmhp motif by solving the steady states algebraically. The exact analytical solution revealed that the transitions of inactivated Rtg2p stem from the molecular titration. Furthermore, the competitive binding between Rtg2p and Bmhp with Mksp forms a binary switch that could process the analog input signal into digital information.

### Frequency modulation of mitochondrial retrograde signaling

In addition to the dose response, we further investigated the frequency modulation of the RTG signaling. Bmh/Mksp heterodimer, and nuclear Rtg1p, and Rtg3p in response to a sinusoidal damage signal of multiple frequencies exhibited low-pass filter behavior (Fig 4 and Fig S4). The recovery rate of Bmh/Mksp synthesis was slower than its degradation, leading to a sign-sensitive delay that keeps the switch on after damage is detected [37], while Bmh/Mksp can be slowly synthesized for a long period of approximately 3 × 10^3^ s compared to the transition of less than 10 s. Although the concentration of Bmh/Mksp recovers slowly after stimulation, Rtg1p and Rtg3p are still able to convey input signal dynamics. At a low frequency, both reach high and low saturation states in a sinusoidal input cycle and produce a square wave pattern. As the frequency becomes high, the amplitude of output diminishes with drifting, indicating that mitochondrial damage signal is low-pass filtered by retrograde signaling. Moreover, the delay of RTG translocation may contribute to the instability and compromise robustness. To investigate the feedback instability of mitochondrial retrograde signaling, we further derived a Bode plot of the retrograde response in the stationary stage (Fig S4). In the frequency response of Bmh/Mksp, the gain decreased in two stages separated by 10^−0.5^ Hz. The gain of the high-frequency stage decreased more rapidly than in low frequency stage. On the other hand, while the phase delay increased significantly in the high-frequency region. The cutoff frequency of nuclear Rtg1p and Rtg3p was approximately 10^−2^ Hz as low-pass filters. For nuclear Rtg3p, there was a second cutoff frequency at approximately 10^−0.5^ Hz with a more rigid downhill slope. Finally, the Nyquist stability theorem was applied to identify the closed-loop stability of mitochondrial retrograde signaling [38]. Despite the phase delay, retrograde signaling remained stable in response to the damage signal. The phase and gain margin derived from the Bode plots can further indicate the stability tolerance to delay caused by the molecular communication like protein translocation. The low-pass filtering can remove the perturbation that is usually in high frequency (Fig 4a) [39].

**Fig 4.**
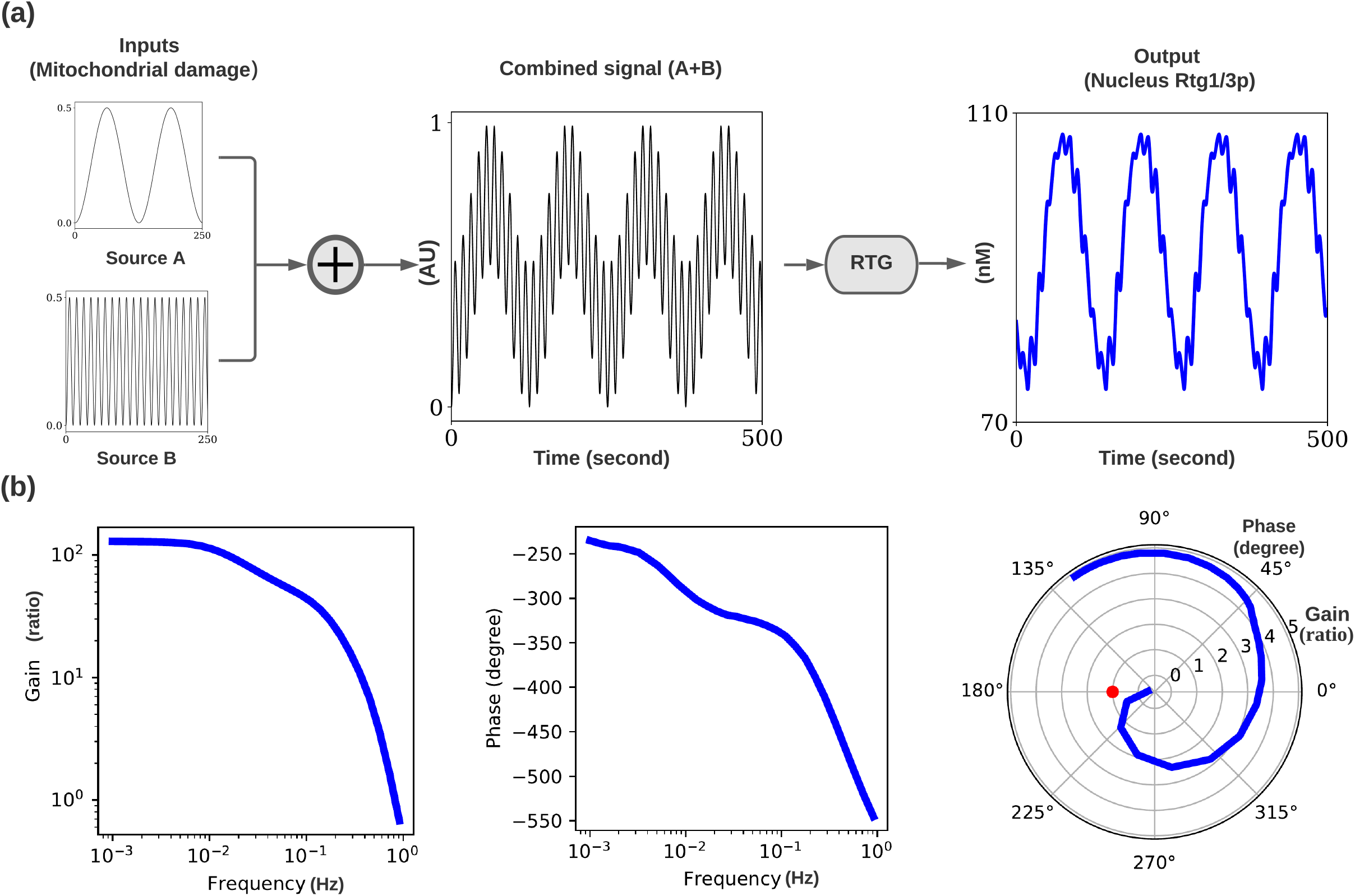
Frequency modulation of RTG pathway. (a) Multiplex processing of RTG signaling. The multiplexed signals from two individual mitochondria are low-pass filtered and delayed with respect to the frequency. The two inputs are set at 0.05 rad/s (source A) and 0.5 rad/s (source B) with the same phase. The overall effect on Rtg2p activation is summed with equal weights, and the combined signal is further modulated by the RTG pathway and detected by nucleus Rtg1/3p as the output. (b) Frequency response to mitochondrial damage, Rtg3p as an example. (Left) Gain plot. Low-pass filtering is observed by screening the input frequency from 10^−3^ to 1 Hz with sinusoidal wave. The gain of nucleus Rtg3p is defined as the ratio of output to input amplitude measured by the peak-to-peak distance. (Middle) Phase plot. The delay of output is increased by the input frequency. With the time deviation between the peaks of output and input, the delayed Rtg3p signal is described by degrees in respect of the input frequency. (Right) The Nyquist plot, combined with the gain and phase plot, reveals the closed-loop stability of the proposed model. The radius represents the gain (dB), and the angle labels the degree of phase. The red dot represents the critical point (−1 + 0*i*). The proposed model is stable if the Nyquist curve excludes the critical point.

### The intensity of the mitochondrial damage signal decreases the robustness of Rtg1/3p translocation

Though the dose-response curve describes the communication between mitochondria and nucleus in a deterministic point of view, the molecular noise causes uncertainty and further limits the information conveyed to the nucleus [62, 63]. To understand how noise influences robustness along with the damage signal, the ODE model was further extended to Chemical Langevin Equations [64]. The proposed deterministic model is incorporated with the random chemical reaction, and the potential landscape of nuclear Rtg1/Rtg3p was derived by screening the damage input with ensemble simulations (Fig 5 and Method Details.). Along with the damage signal, the sigmoid curve of the potential landscape was consistent with the deterministic results of Rtg1/Rtg3p (Fig 5b). Further, three conjugated steady states were identified in response to a constant damage (Fig 5). A steady state represents the Rtg3p distribution that is in high probability. The conjugated steady states became distanced as the input signal increased, indicating that noise was amplified and caused an increased bandwidth, leading to a compromised signal-to-noise ratio. Notably, three conjugated steady states were further investigated by measuring the output distribution for a given input (Fig 5). The three conjugated steady states were identified by the peak of the probability, which represented the high likelihood of the Rtg1/Rtg3p concentration in a given input. The conjugated steady states were located within the bandwidth of the probability density function (PDF), and the length of bandwidth was increased by the signal. This result suggested that mitochondrial retrograde signaling was not only bistable but also a toggle-switch with multiple locally conjugated stable steady states (Fig 5c). In addition, the uncertainty of the output was worsened by a high input signal, which might compromise information relayed between mitochondria and the nucleus when the former continues to send high-intensity messages.

**Fig 5.**
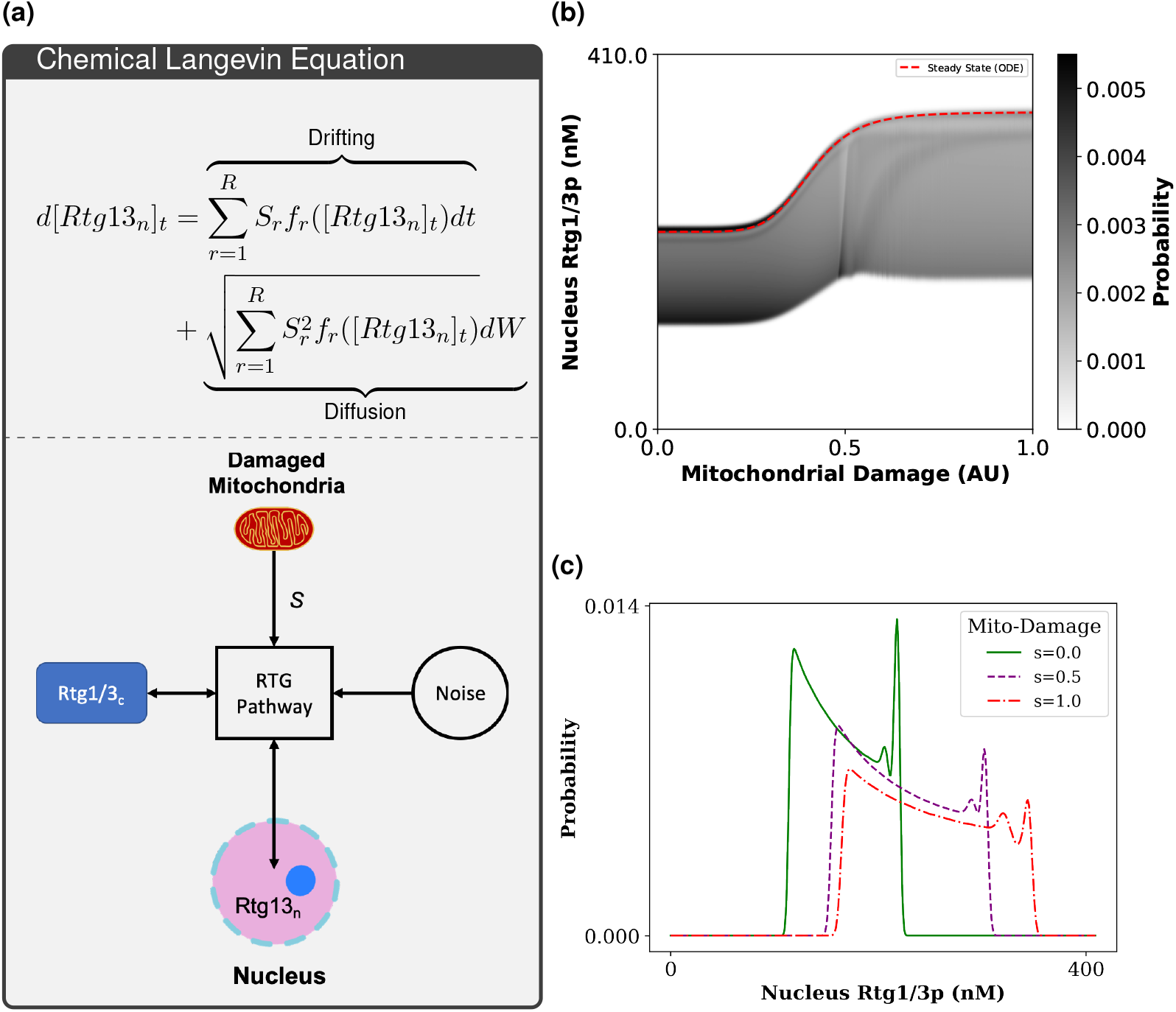
Robustness of Rtg1/3p translocation in response to mitochondrial damage signal. (a) The potential landscape of the nuclear Rtg1/Rtg3p response to mitochondrial damage signals. The sigmoid transition is identified in Fig S3a, however, the bandwidth (the dark area in the vertical section) increases in parallel to the damage signal. The damage signal is sampled in 100 grids and assigned to the Chemical Langevin Model in order to simulate the stochastic process. Every combination of parameters is applied with 100-time length (arbitrary unit) with automatic time step. The y-axis is limited within the range of 0 and the maximum concentration of Rtg3p. A heatmap shows the probability with a color bar on the right side. (See Method Details.) (b) The probability density function (PDF) with respect to Rtg1/Rtg3p influenced by damage signal. The distribution is simulated by setting the input signal as 0 (blue, minimal input), 0.45 (orange, intermediate input), and 1 (green, maximum input). The PDF is measured by time series simulation and estimates the probability via kernel density estimation

## Discussion

The RTG retrograde signaling pathway is as an important mechanism for mitochondrial quality control as it serves as both a sensor and a reporter of the mitochondrial health status [13]. Quantitative and systemic analysis of this major communication channel will help us elucidate the crosstalk between mitochondria and the nucleus, which maintains cellular function under physiological and pathological conditions. Herein, we constructed a novel mathematical model of mitochondrial retrograde signaling in yeast, applying microscopic data of the RTG pathway knockout experiments with gene and protein expression data for model validation. The differential equation-based model provides a comprehensive approach for studying the dynamic response of mitochondrial retrograde signaling, which has been experimentally challenging due to the lack of means for precise mitochondrial quality manipulation in real time. Retrograde signaling dynamics were observed after introducing the input signal with step and sinusoidal waveforms. Moreover, we applied an analytical solution to identify the source of ultrasensitivity in the degradation of the Bmh/Mksp heterodimer.

The switch-like behavior of the yeast RTG pathway was previously described by [12]. We validated switch-like behavior in this retrograde signaling model (Fig 1) through mathematical formulation and numerical simulation. We observed that switch-like behavior resulted from the competitive binding between active Rtg2p and Bmhp. Besides, the effect of molecular titration triggered a two-staged transition in inactive Rtg2p. Ultrasensitivity is typical in cellular communication, allowing cells to remove noise and amplify input signals [40–42]. Rtg2p senses mitochondrial quality through the modulation of Bmhp and Mksp, facilitating detection via competitive binding. The “all-or-none” response can further filter the fluctuation and adjust to the unpredictable environment.

The current study further elaborates on the properties of signal modulation via Rtg1/3p translocation following static and sinusoidal damage signals. Rtg1p acts as both a positive and negative regulator [18], retaining the transcription factor Rtg3p in the cytoplasm. This considerably constrains the valid parameter set. Further mechanistic details were identified by the proposed model: the two-staged transition of Rtg3p translocation in step response and the sign-sensitive delay of Rtg3p with a short ON delay in oscillation signal. Without NLS modification, Rtg1p is required to retain Rtg3p in the cytoplasm when a mitochondrial damage signal is absent, which is one of the criteria of our model. Rtg1p exerts an anchoring effect on Rtg3p by forming a heterodimer, which may not permeate through the nuclear membrane. The translocation of Rtg1p is not affected by the sign of the input change and reveals a simple diffusion process with a capacity-charging pattern. The frequency response further revealed the low-pass filtering of the RTG pathway, which can reject high-frequency fluctuations and retain the overall information of the input signal for requesting nuclear supply (Fig 6).

**Fig 6.**
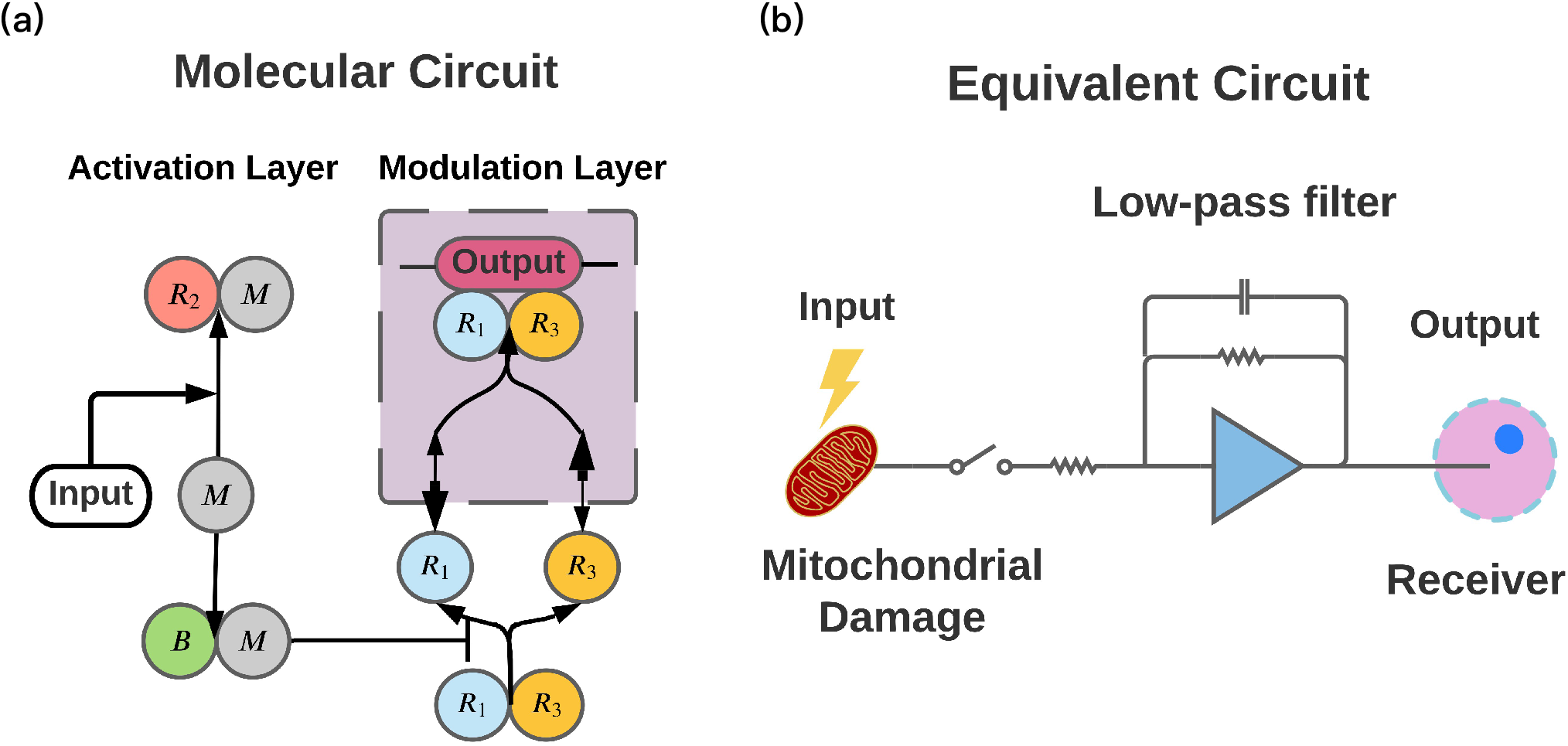
Mitochondria retrograde circuit. (a) Mitochondrial retrograde signaling facilitates communication between the mitochondria and nucleus. The retrograde molecular circuit transduces the signal through the activation layer and the modulation layer to sense and to report the signal to the nucleus. (b) Retrograde signaling circuit was switched on by a mitochondrial damage signal, while a low-pass filter maintained the robustness of the pathway.

In the current study, we demonstrated feedback stability of RTG signaling using Bode and Nyquist diagrams [38]; the communication between mitochondria and the nucleus is a two-way process [43]. The phase and gain margin in the Bode plot determined the stable region when cascading with these functional units under mitochondrial oscillation.

Our model also elucidates how the RTG pathway processes the input signal as a low-pass filter, reporting status to the nuclear genome, which may, in turn, support the synchronization of mitochondrial membrane potential in yeast [44–46] and insulin secretion from pancreatic cells [47]. Moreover, mitochondria undergo energization cycles driven by the ultradian clock, contributing to respiratory oscillations of mitochondria [44–46]. These oscillations may be a source of reactive oxygen species generated by mitochondria [45], and further interfere the retrograde signaling with sinusoidal input.

The mitochondrion acts as a biochemical signaling hub in eukaryotic cells [48]. Apart from energy production, this semi-autonomous organelle also participates in various metabolic reactions related to aging [49–51], cellular communication [4, 47] and cell viability [26, 30]. The RTG response facilitates the adaptation of eukaryotic organisms to unfavorable factors [26, 27]. In yeast, the key role of RTG signaling is the regulation of mitochondrial function and metabolic reprogramming, including the maintenance of intracellular glutamate supplies [27, 28]. Studies have reported the association between the retrograde response and drug resistance in yeast [27, 29, 30]. For example, Rtg3p has been reported as essential for antifungal drug tolerance associated with the formation of the fungal plasma membrane, which contains ergosterol [31]. The Δ*rtg3* mutant of pathogenic yeast, *Candida albicans*, exhibits compromised biofilm formation and epithelial cell adherence, resulting in reduced infectivity of nematode *modelCaenorhabditis elegans* [29]. Moreover, in *Saccharomyces cerevisiae*, pleiotropic drug resistance is activated by RTG signaling, inducing the transcription of pleiotropic drug resistance genes [27]. RTG signaling is also associated with aging, highlighting yeast as a model organism for the study of mitochondrial quality control and its influences on lifespan [22].

Retrograde signaling is connected to glutamate synthesis, mTOR signaling, and other feedback circuits [5, 13, 52]. The RTG pathway is a major factor in cellular adaptation, serving a cytoprotective role via metabolic reprogramming [6, 27]. The robustness of retrograde signaling is essential for adjusting to unfavorable conditions [26]. For instance, RTG signaling contributes to drug resistance in pathogenic yeast, such as *C. albicans*, which is responsible for approximately 80% of major systemic fungal infections [27, 53]. Various antifungal drugs target mitochondria-related amino acid and ergosterol metabolism [54,55], and the RTG pathway enables pathogenic yeast to overcome therapeutic agents by sensing the loss of mitochondrial functionality and subsequently reprogramming their metabolism [27, 31, 56]. Hence, the yeast retrograde signaling system represents a target for the development of novel antifungal drugs [27]. However, how yeast senses mitochondrial stress via the RTG pathway in a dynamic environment remains unclear.

In this study, we applied mathematical modeling to unravel the dynamics of RTG signaling from a control system point of view. We further identified the threshold of the RTG response, which can be used for determining the optimal dosages for the elimination of pathogenic yeast without triggering RTG signaling. The frequency response reveals how yeast manages to resist fluctuation via low-pass filtering of the RTG pathway, as well as the hysteresis effect, which maintains retrograde signaling as the damage signal fluctuates, indicating that persistent treatment with a concentration slightly below the RTG response threshold is an optimal strategy against pathogenic yeast. Therefore, comprehensively understanding the qualitative properties of retrograde signaling may shed light on signaling mechanisms and provide potential therapeutic strategies.

## Acknowledgment

The work was supported by the Ministry of Science and Technology in Taiwan (MOST-109-2636-B- 002 -001 and MOST-110-2636-B- 002 -001 grants to AW).

## Data and Code Availability

The source code is available at https://github.com/NTUMitoLab/MitoRetroDynamics. Julia toolbox developed for potential map generator, ultrasensitivity analysis, and steady state explorer are also available at GitHub.

## Method Details

### Boolean model of mitochondrial retrograde signaling

To summarize the systematic behavior of mitochondrial retrograde signaling, a Boolean model was established to discretize the mechanism of protein translocation associated with mitochondrial damage. In this model, The population of *N* mitochondria is indexed into a vector 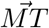(Eq. 1).

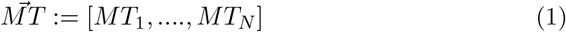

Though it is known that damaged mitochondria activate Rtg2p and promote the dephosphorylation of Rtg3p by decreasing the Bmh/Mksp heterodimer [16, 18], it is unclear how Rtg2p senses mitochondrial dysfunction. Therefore, we assume mitochondria activates Rtg2p simultaneously under Michaelis-Menten kinetics, and the activation rate of Rtg2p is influenced by the weighted linear summation of mitochondrial damage signals (*S*) with their volume(Eq. 2).

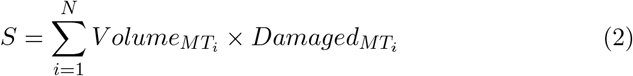

In the Boolean model, the signal (*S*) is set to 0 for the healthy mitochondrial population and to 1 when mitochondria are damaged, as in *ρ*^0^ yeast cells which have no mitochondrial DNA. These genetically damaged mitochondria have no respiratory functionality and constitutively activate retrograde signaling, which causes the translocation of Rtg1p and Rtg3p, when cells contain intact RTG components (Table S1). The translocation of Rtg1p and Rtg3p can switch on the retrograde response, including the upregulation of *CIT2* [16].

Sekito et al. [18] determined the nuclear concentration of Rtg1p, Rtg2p and Rtg3p relative to cytoplasmic levels via GFP labeling, RTG knockouts, and the comparison between *ρ*^+^ and *ρ*^0^ cells. Therefore, the translocation event, existence of RTG components, and mitochondrial functionality can be summarized into a Boolean table, describing the system behavior of the RTG signaling pathway (Table 1) [18]. The Boolean relation between mitochondrial damage (input) and the translocation of Rtg1p or Rtg3p can be further simplified by Karnaugh map (Fig 2a).

### From Boolean model to an ordinary differential equation-based model

To investigate the dynamics of mitochondria-nucleus communication, an ordinary differential equation-based model with two compartments (cytosol and nucleus) was developed to simulate RTG activation and translocation (Fig 1). Reactions of mitochondrial retrograde response were derived based on the law of mass action and Michaelis-Menten kinetics. There are total of 17 differential equations(S1.1) and 24 kinetic coefficients (Table S3) to describe the protein dynamics with different activation states or locations in the RTG signaling pathway.

The activation layer consists of Rtg2p, Mksp and Bmhp. This layer is constantly suppressed by Bmhp-Mksp heterodimer, and conveys the mitochondrial damage signal via Rtg2p activation. Later on, Rtg1p and Rtg3p form the modulation layer. Rtg3p contains nuclear localization sequence (NLS) and is poly-phosphorylated when the RTG pathway is inactive. In contrast, upon mitochondrial damage, Rtg3p is less phosphorylated and tends to translocate into the nucleus. The process of transcription factor translocation of modeled with simplified mass kinetics (Eq. 3) as previously described [59].

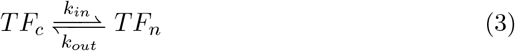

where *TF*_∗_ represents transcription factor, and the subscripts ∗ describes the location with *n* representing nucleus and *c* the cytosol. The compartmental model is further performed with the following equalities:

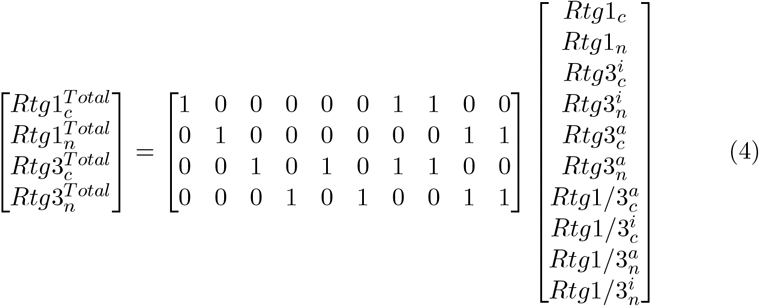

where 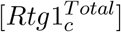 is defined as the total cytoplasmic concentration of Rtg1p and heterodimers containing Rtg1p. The localization is labeled with cytosol (*c*) and nucleus (*n*) in subscript. *a* and *i* in superscript represent partial phosphorylation and hyper-phosphorylation forms of Rtg3p. The dephosphorylation process turns on the nucleus localization sequence (NLS) in Rtg3p [18].

### The differential equation-based model of RTG signaling

A mathematical model was proposed to fulfilled the current understanding in signaling RTG pathway as summarized previously [18, 32]. The RTG pathway model was based on differential equations to explore the system dynamics. The model was composed of biochemical reaction, Michaelis-Menten enzymatic kinetics, and Hill equation. The activation layer conveys the mitochondrial damage signal via Rtg2p activation. The modulation layer translocates Rtg3p and Rtg1p to nucleus to initiate RTG response.

1. **Activation layer** : Rtg2p and mitochondrial damage signal (*s*).

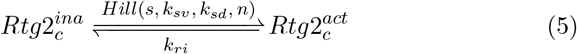

where 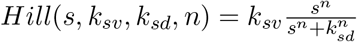. Competitive interaction between Rtg2p and Bmhp on Mksp.

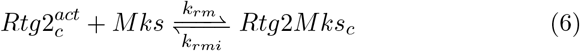

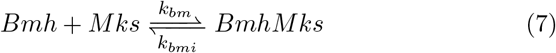
2. **Modulation layer**: Translocation of Rtg3p, Rtg1p and their derivatives [18, 33]. The constant inhibition of Bmh/Mksp is described by

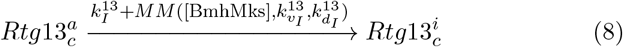

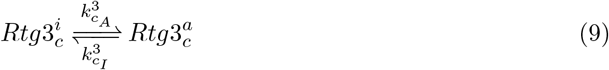

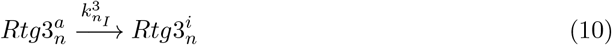

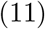

where 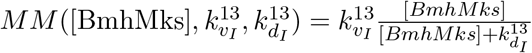 The formation of Rtg1/3

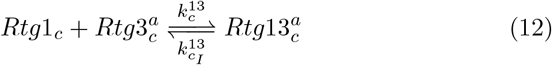

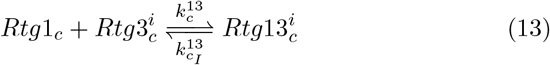

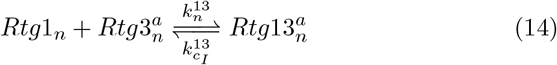

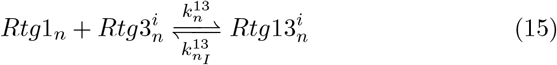

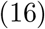 Translocation of Rtg1/3p

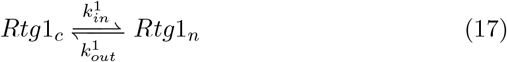

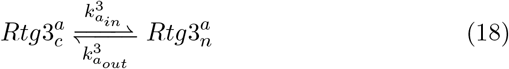

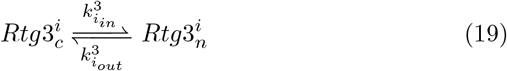

To determine the relative concentrations of proteins in the RTG pathway, the expression level of RTG genes under normal and osmotic stress conditions (GEO access number: GSE59659) [35], and protein expression level obtained from *Saccharomyces cerevisiae* fusion library [34] were analyzed and used for determining the total concentration of RTG proteins(Fig S1).

The differential equations are described in Section S1.1. The parameter searching algorithm is in Section S1.2, and the final parameter set is listed in Table S3.

### Parameter searching with the qualitative data of protein translocation

The parameters determined the steady-state behavior with respect to mitochondrial damage. A valid parameter set should make the model fulfill all conditions observed in [18] and [28] (Table 1). The parameter set was sampled in log-uniform with a fixed domain, and the relative amount of translocation kinetic coefficients is considered. To elaborate, the inward coefficient of nuclear Rtg3p with an active 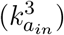 was higher than the inactive one 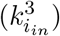. Conversely, 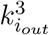 was higher than 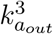 [59].

The initial values of each component were sampled based on the RNA expression levels. The relative amount of protein under both normal and stress conditions was considered. The Boolean relations were tested by setting deleted proteins near zero, including modifications and complexes.

Once the parameters were sampled, the steady state of the model was derived by a dynamic solver, which provides an automatic switch between stiff and nonstiff algorithms when solving the differential equations. Furthermore, we chose Order 2/3 L-Stable Rosenbrock-W method as the stiff solver, and Tsitouras 5/4 Runge-Kutta method for the nonstiff section [60].

The unstable solutions were discarded if the iteration exceeded 10^6^ with a deviation of derivatives beyond 10^−8^ in total. Further, the ratio of the nuclear concentration to cytoplasmic one was used to determine protein translocation. A nucleus-to-cytosol ratio larger than 1.5 was defined as a translocation event and vice versa. The translocation process under each deletion condition was compared with the Boolean model.

The differential equations were solved by DifferentialEquations.jl (https://diffeq.sciml.ai/stable/), a open-source package for scientific computing implemented in the Julia programming language (https://julialang.org/). The parameter searching was performed on one machine with 30 Intel Xeon Processors in TWCC cloud service (https://www.twcc.ai/) for 10^9^ iterations.

### The analytical solution of the activation layer

The acticvation layer contains three cytoplasmic proteins: Rtg2p, Bmh1/2p and Mks1p [13, 16, 21]. A decline in mitochondrial membrane potential (ΔΨ_*m*_) leads to activation of Rtg2p, and interferences with the formation of Mks1p-Bmh1/2p complex, thereby impeding the nucleus accumulation of Rtg3p by inhibiting its partial dephosphorylation [16].

The reaction is summarized in the form of competitive binding with three components. Rtg2p and Bmhp (including Bmh1p and Bmh2p) are two ligands that compete for the Mks1p (Eq.22). The input of the activation layer is the mitochondrial damage (*s* in Eq.22) which can be the decline of its ΔΨ_*m*_, and further defined in the domain between 0 and 1 which represents the continuous state from healthy to the damaged. Further, the activation of Rtg2p induced by mitochondrial damage is modeled by a Hill equation. In the proposed model, the kinetic coefficients and total concentration of each protein are regarded as constants (Eqs. 25 and 28).

With the verified parameter set, the two-staged transition of Rtg2p activation induced by the mitochondrial damage signal was identified. The first stage of transition is a sigmoidal curve with the following steady region as the damage signal turns high, revealing the behavior of ultrasensitivity. After a certain value, the active form of Rtg2p increases in a convex trend with a slower slope compared to the first transition. Furthermore, by screening possible Hill coefficient of Rtg2p activation (*n^s^*), we found that the threshold of the first transition, rather than the stiffness of the curve, becoming larger when *n* increases.

To further understand these phenomena, it is necessary to derive the analytical solution of steady states to understand the input-output relation of mitochondrial damage and the total concentration of Bmh/Mksp heterodimer. In the following paragraph, we used 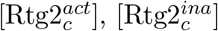, [Bmh] and [Mks] to represent their steady-state concentration.

### The competitive binding of Bmhp and Rtg2p with Mksp is the source of the ultrasensitivity

The modified competitive binding model from [61].

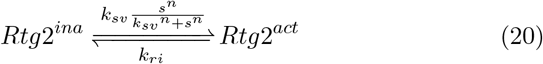

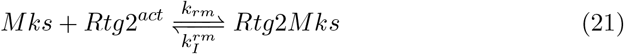

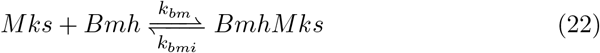

The dissociation coefficients are defined as

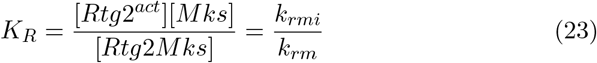

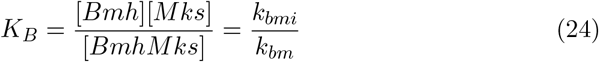

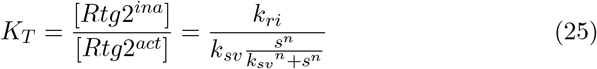

where *K_R_* and *K_B_* are the dissociation coefficients for the Mksp’s binding of Rtg2p^*act*^ and Bmhp. For the Rtg2p activation, We assume the input influcences the activation of Rtg2p^*ina*^ in the manner of Hill formula (Eq.20 and Eq.25). To avoid zero division, we set the domain of input as *s* ∈ (0, 1].

The conservation rule requires that

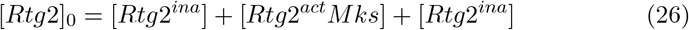

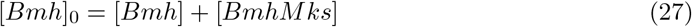

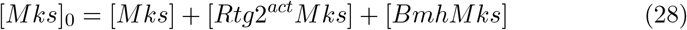

where 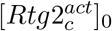, [*Bmh*]_0_ and [*P*_0_] represent total concentration, and are assumed constants. Therefore, we can make [*Bmh*] and [*Rtg2^act^Mks*] into the dependent variables of [*Mks*] and their total concentration.

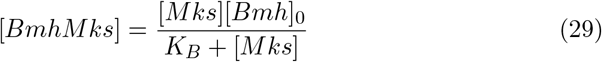

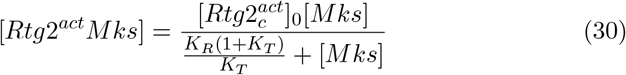

To derive the polynomial function of [*Mks*], we can put Eq.29 and Eq.30 together:

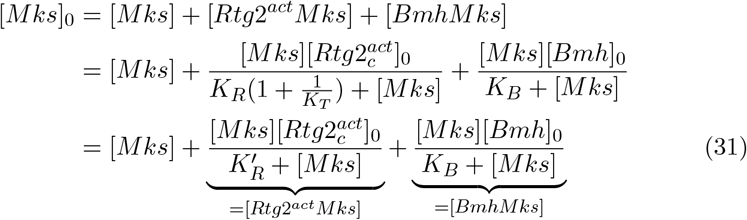

where 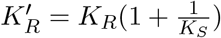 is the dissociation coefficient influenced by the input coefficient *K_T_*(Eq.25).

Finally, we can get the cubic polynomial equation of [*Mks*]from Eq.31:

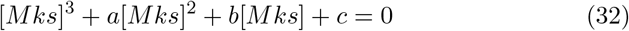

where

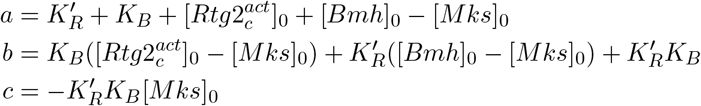

The polynomial equation of [*Mks*] can be solved by applying trigonometry [61],

The analytical solution of competitive ligand binding

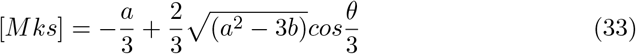

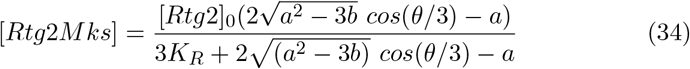

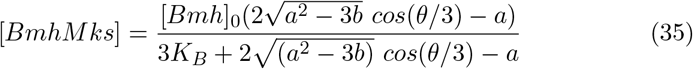

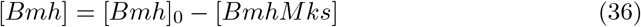

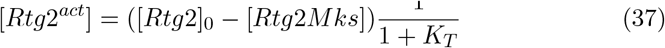

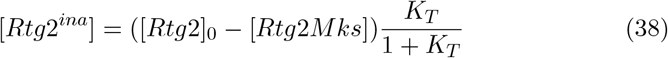

where 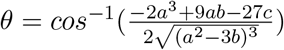.

### Analysis ultrasensitivity with relative amplification approach

The relative amplification approach is applied to quantify the local and global sensitivity by estimating the equivalent Hill coefficient [36]. When the response curve starts with a high basal level, this approach provides a more accurate approximation compared to the method using EC90:EC10 ratio [36, 57]. The response coefficient (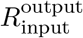, Eq 39) is used to quantify the relative change in response to the stimulus.

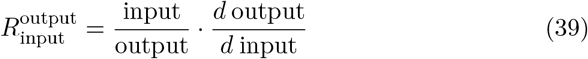

Furthermore, to circumvent the deviation from a high basal level, the activated fraction (*f*, Eq 40) is defined as the activated ratio of output between the maximum and basal response [36].

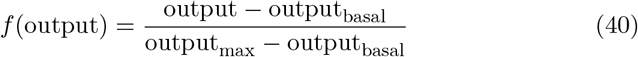

where *f* ∈ [0, 1], output is the function of input. Besides, output_max_ and output_basal_ represents the maximum and minimum of outputs respectively, which makes activated fraction (*f*) independent from the basal activation.

The advantage of applying the response coefficient (*R*) and activated fraction (*f*) is that the classical Hill equation can be converted to a linear equation (Eq 41) [36].

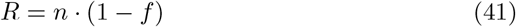

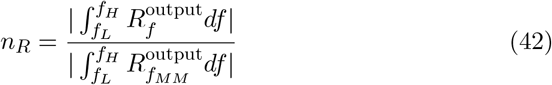

where *f_H_* and *f_L_* are the maximum and minimum of activation fraction in a given region.

### Frequency response of RTG signaling

To measure the RTG response to the dynamical input, the original ODE model is converted into non-autonomous differential equations with mitochondrial damage signal (*s*) as input (Eq.43).

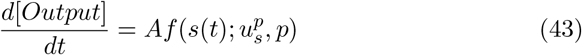

where *A* is the output map

Furthermore, the sinusoidal function with frequency between 10^−3^ Hz and 1 Hz was used to conduct the frequency analysis to reveal the break point in Bode diagram. In this study, we focus on the concentrations of Bmh/Mksp heterodimer and Rtg1/3p nucleus accumulation in response to the sinusoidal input.

The nonautonomous model is solved by Rosenbrock23 with automatic switching for Tsit5 solver. The Rosenbrock23 is a stiff ODE solver which can deal with more general situation than Tsit5, a non-stiff solver, but with lower speed [60]. By the aid of automatic switching stiff/non-stiff method of DifferentialEquations.jl [60], the simulation process can be significantly sped up.

### Stochastic simulation of Rtg1/3p translocation in response to mitochondrial damage

The stochastic model is modified from the ODEs with the theory of Chemical Langevin Equation (CLE) [64]. The derivative of nucleus Rtg1/3p is composed of the drifting term from the deterministic model, and the diffusion term caused by the Brownian motion. The process is described with the stochastic differential equation in Eq 44.

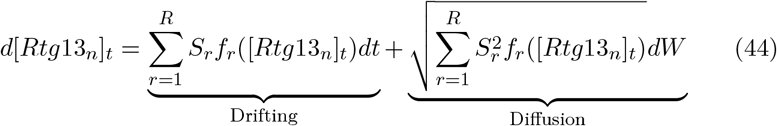

where *f_r_* is the propensity of *r^th^* reaction, *S_r_* the net change of the reaction *r*,and *dW* represents the Wiener process [65].

The CLE was solved using the Euler-Maruyama method with adaptive time stepping, and simulated for 100 seconds. Further, the time series simulation was used to derive the likelihood of the output state with kernel density estimation as illustrated in Fig 5.

