## Supplement for "Mathematical Modeling and Analysis of Mitochondrial Retrograde Signaling Dynamics"

### Supporting Information for Mathematical Modeling and Analysis of Mitochondrial Retrograde Signaling Dynamics

### S1 The Structure of the Differential Equation-Based Model of Mitochondrial Retrograde signaling in Yeast

#### S1.1 Differential equation-based model

Activation Layer

$$\frac{ds(t)}{dt} = 0 \tag{S1}$$

$$\frac{dRtg2_c^i}{dt} = -\frac{k_{sv}\,(s\,(t))^{n_s}}{{k_{sd}}^{n_s} + (s\,(t))^{n_s}} \mathrm{Rtg2}_\mathrm{c}^\mathrm{ina}\,(t) + k_I^r \mathrm{Rtg2}_\mathrm{c}^\mathrm{act}\,(t) \tag{S2}$$

$$\frac{dRtg2_{act\_c(t)}}{dt} = \frac{k_{sv}\,(s\,(t))^{n_s}}{{k_{sd}}^{n_s} + (s\,(t))^{n_s}} \mathrm{Rtg2}_\mathrm{c}^\mathrm{ina}\,(t) - k_I^r \mathrm{Rtg2}_\mathrm{c}^\mathrm{act}\,(t) - k^{rm} \mathrm{Rtg2}_\mathrm{c}^\mathrm{act}\,(t)\,\mathrm{Mks}\,(t) + k_I^{rm} \mathrm{Rtg2Mks_c}\,(t) \tag{S3}$$

$$\frac{dMks(t)}{dt} = -k^{rm} \mathrm{Rtg2}_\mathrm{c}^\mathrm{act}\,(t)\,\mathrm{Mks}\,(t) + k_I^{rm} \mathrm{Rtg2Mks_c}\,(t) - k^{bm} \mathrm{Bmh}\,(t)\,\mathrm{Mks}\,(t) + k_I^{bm} \mathrm{BmhMks}\,(t) \tag{S4}$$

$$\frac{dRtg2Mks_{c(t)}}{dt} = k^{rm} \mathrm{Rtg2}_\mathrm{c}^\mathrm{act}\,(t)\,\mathrm{Mks}\,(t) - k_I^{rm} \mathrm{Rtg2Mks_c}\,(t) \tag{S5}$$

$$\frac{dBmh(t)}{dt} = -k^{bm} \mathrm{Bmh}\,(t)\,\mathrm{Mks}\,(t) + k_I^{bm} \mathrm{BmhMks}\,(t) \tag{S6}$$

$$\frac{dBmhMks(t)}{dt} = k^{bm} \mathrm{Bmh}\,(t)\,\mathrm{Mks}\,(t) \tag{S7}$$

$$- k_I^{bm} \mathrm{BmhMks}\,(t) \tag{S8}$$

#### Modulation Layer

$$\frac{dRtg13_{a\_c(t)}}{dt} = -\left(k_I^{13} + \frac{Rtg2_c^{ina} k_{v_I}^{13} \mathrm{BmhMks}(t)}{k_{ID}^{13} + \mathrm{BmhMks}(t)}\right) \mathrm{Rtg13_c^a}(t) \tag{S9}$$

$$+ k_c^{13} \mathrm{Rtg1_c}(t)\, \mathrm{Rtg3_c^a}(t) - k_{c_I}^{13} \mathrm{Rtg13_c^a}(t) \tag{S10}$$

$$\frac{dRtg13_{i\_c(t)}}{dt} = \left(k_I^{13} + \frac{Rtg2_c^{ina} k_{v_I}^{13} \mathrm{BmhMks}(t)}{k_{ID}^{13} + \mathrm{BmhMks}(t)}\right) \mathrm{Rtg13_c^a}(t) \tag{S11}$$

$$+ k_c^{13} \mathrm{Rtg1_c}(t)\, \mathrm{Rtg3_c^i}(t) - k_{c_I}^{13} \mathrm{Rtg13_c^i}(t) \tag{S12}$$

$$\begin{aligned}\frac{dRtg3_{i\_c(t)}}{dt} =& - k_{c_A}^3 \mathrm{Rtg3_c^i}(t) + k_{c_I}^3 \mathrm{Rtg3_c^a}(t) \\ & - k_c^{13} \mathrm{Rtg1_c}(t)\, \mathrm{Rtg3_c^i}(t) + k_{c_I}^{13} \mathrm{Rtg13_c^i}(t) \\ & - k_{i_{in}}^3 \mathrm{Rtg3_c^i}(t) + k_{i_{out}}^3 \mathrm{Rtg3_n^i}(t)\end{aligned}$$

$$\begin{aligned}\frac{dRtg3_{a\_c(t)}}{dt} =& k_{c_A}^3 \mathrm{Rtg3_c^i}(t) - k_{c_I}^3 \mathrm{Rtg3_c^a}(t) \\ & - k_c^{13} \mathrm{Rtg1_c}(t)\, \mathrm{Rtg3_c^a}(t) + k_{c_I}^{13} \mathrm{Rtg13_c^a}(t) \\ & - k_{a_{in}}^3 \mathrm{Rtg3_c^a}(t) + k_{a_{out}}^3 \mathrm{Rtg3_n^a}(t)\end{aligned} \tag{S13}$$

$$\begin{aligned}\frac{dRtg3_{a\_n(t)}}{dt} =& - k_{n_I}^3 \mathrm{Rtg3_n^a}(t) - k_n^{13} \mathrm{Rtg1_n}(t)\, \mathrm{Rtg3_n^a}(t) \\ & + k_{n_I}^{13} \mathrm{Rtg13_n^a}(t) + k_{a_{in}}^3 \mathrm{Rtg3_c^a}(t) \\ & - k_{a_{out}}^3 \mathrm{Rtg3_n^a}(t)\end{aligned} \tag{S14}$$

$$\begin{aligned}\frac{dRtg3_{i\_n(t)}}{dt} =& k_{n_I}^3 \mathrm{Rtg3_n^a}(t) - k_n^{13} \mathrm{Rtg1_n}(t)\, \mathrm{Rtg3_n^i}(t) \\ & + k_{n_I}^{13} \mathrm{Rtg13_n^i}(t) + k_{i_{in}}^3 \mathrm{Rtg3_c^i}(t) \\ & - k_{i_{out}}^3 \mathrm{Rtg3_n^i}(t)\end{aligned} \tag{S15}$$

$$\begin{aligned}\frac{dRtg1_{c(t)}}{dt} =& - k_c^{13} \mathrm{Rtg1_c}(t)\, \mathrm{Rtg3_c^a}(t) \\ & + k_{c_I}^{13} \mathrm{Rtg13_c^a}(t) - k_c^{13} \mathrm{Rtg1_c}(t)\, \mathrm{Rtg3_c^i}(t) \\ & + k_{c_I}^{13} \mathrm{Rtg13_c^i}(t) \\ & - k_{in}^1 \mathrm{Rtg1_c}(t) + k_{out}^1 \mathrm{Rtg1_n}(t)\end{aligned} \tag{S16}$$

$$\tag{S17}$$

$$\begin{aligned}\frac{dRtg1_{n(t)}}{dt} =& - k_n^{13} \mathrm{Rtg1_n}(t)\, \mathrm{Rtg3_n^a}(t) \\ & + k_{n_I}^{13} \mathrm{Rtg13_n^a}(t) - k_n^{13} \mathrm{Rtg1_n}(t)\, \mathrm{Rtg3_n^i}(t) \\ & + k_{n_I}^{13} \mathrm{Rtg13_n^i}(t) + k_{in}^1 \mathrm{Rtg1_c}(t) - k_{out}^1 \mathrm{Rtg1_n}(t)\end{aligned} \tag{S18}$$

$$\frac{dRtg13_{a\_n(t)}}{dt} = k_n^{13} \mathrm{Rtg1_n}(t)\, \mathrm{Rtg3_n^a}(t) - k_{n_I}^{13} \mathrm{Rtg13_n^a}(t) \tag{S19}$$

$$\frac{dRtg13_{i\_n(t)}}{dt} = k_n^{13} \mathrm{Rtg1_n}(t)\, \mathrm{Rtg3_n^i}(t) - k_{n_I}^{13} \mathrm{Rtg13_n^i}(t) \tag{S20}$$

#### S1.2 Parameter searching with Monte Carlo Simulation

**Algorithm 1:** Framework of finding valid parameter sets of Boolean satisfiability.

```
Data: Boolean relation of deletions and mitochondrial status (test
      conditions in Figure 2.)
Result: Dictionary of parameter set with Boolean satisfication on each
        conditions.
while number of trials ≤ Maximum iteration do
    Sample a parameter array with elements in the range of table;
    Shuffle the order of test conditions;
    Summing up each protein species and redistributed;
    for i ← 0 to number of conditions do
        if condition[i] is missing then
            continue;
        end
        foreach deleted protein j in condition[i] do
            Set protein j ← 0;
        end
        Input ← If Mitochondria are healthy ? 0: 1
        Solve the steady state;
        if Solve failed then
            Break;
        end
        Translocations[i] ← If Nucleus GFP ≥ 1.5× Cytoplasmic GFP ? 1
         : 0
        if Translocation matches with Data then
            Assign valid;
        end
    end
end
```

### S2 Figures and Tables

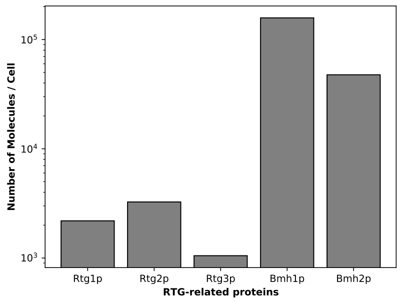

**(a) Expression levels from quantitative Western blot [15]**

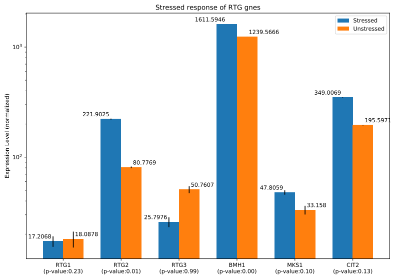

**(b) Differential expressions in response to osmotic stress [16]**

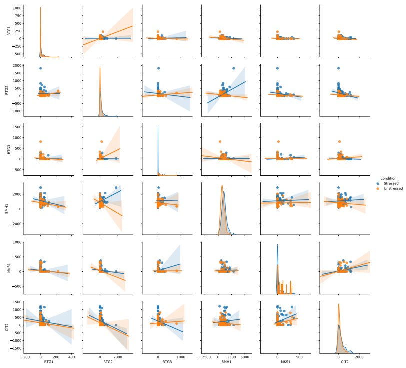

**(c) Correlation of RTG gene expressions in response to osmotic stress [16]**

**Fig S1. Expression levels of RTG proteins.**
(a) The quantitative Western blot data is derived from https://yeastgfp.yeastgenome.org/ [15]
(b-c) The microarray data is derived from the Gene Expression Omnibus(GEO) database (access number GSE59659) [16], and analyzed by D3E algorithm [17].

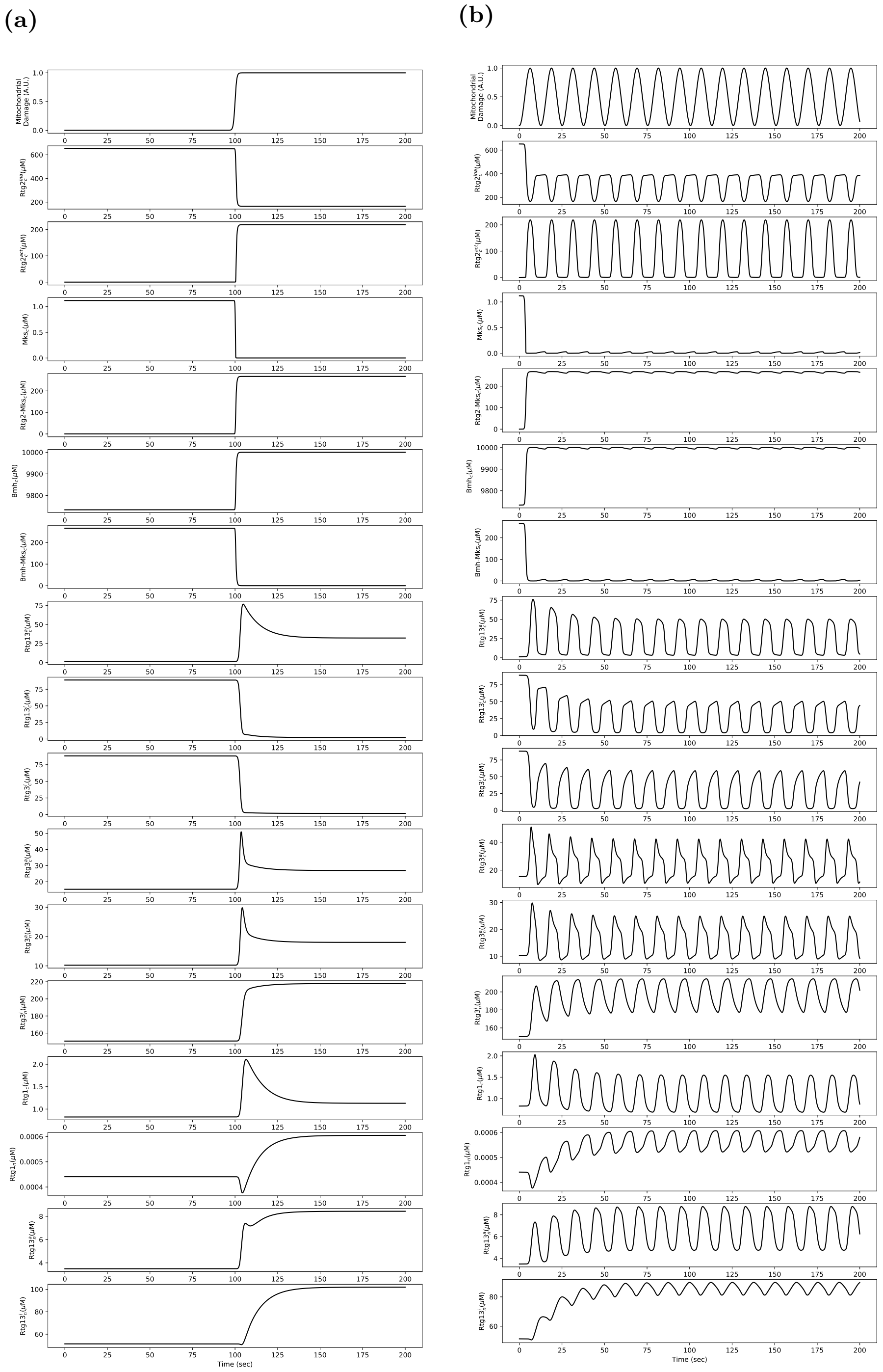

**Fig S2. The retrograde response to mitochondrial damage with sigmoid and sinusoidal waveform.**
(a) When applying sharp sigmoid signal, there is an overshoot effect on Rtg3p$_c^a$, Rtg3p$_n^a$, Rtg1p$_c$, Rtg1p$_c$ and Rtg1/3p$_n^a$. (b) Both waveforms possess similar characteristics: the inactive form of Rtg3p performs with impulse train and Rtg1/3p$_n^a$ with the square wave containing overshoot when the step changing as ON. The Mksp-associated components remain nearly steady with small ripples responded to the periodic input signal.

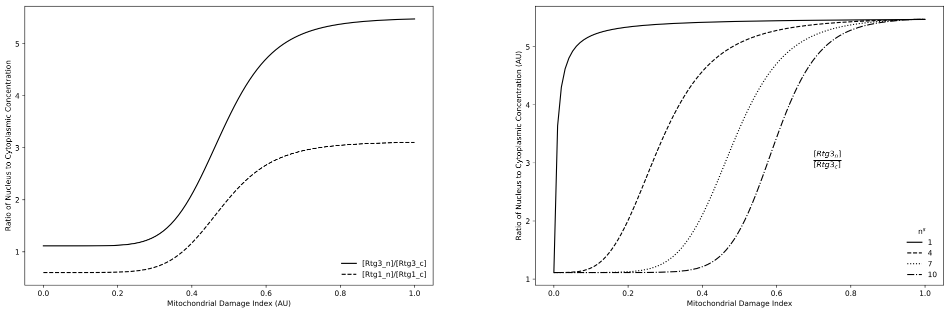

**(a)** **(b)**

**Fig S3. The input-output relation and Hill coefficient.**
(a) The influence of Hill coefficient of input sensing ($n_s$) on nuclear accumulation. The Rtg2p sensing of mitochondrial damage signal is modeled by the Hill equation with the parameter of Hill coefficient, $n^s$. As the mechanism of Rtg2p activation induced by mitochondrial damage remains unclear, the Hill equation is used to generalize the reaction. In our model when the Hill coefficient is equal to 1, the nuclear accumulation is rigidly responding to mitochondrial damage signal. However, when the Hill coefficient is greater than 1, the curve of the phase plot becomes steady, yet gradually increasing the threshold. (b) Phase plot for nuclear accumulation responding to mitochondrial damage signal. Nuclear accumulation is measured by dividing the total nucleus concentration by the total cytoplasmic concentration. Both Rtg1p and Rtg3p accumulate in the nucleus with a monotonous trend when the mitochondrial damage signal is high.

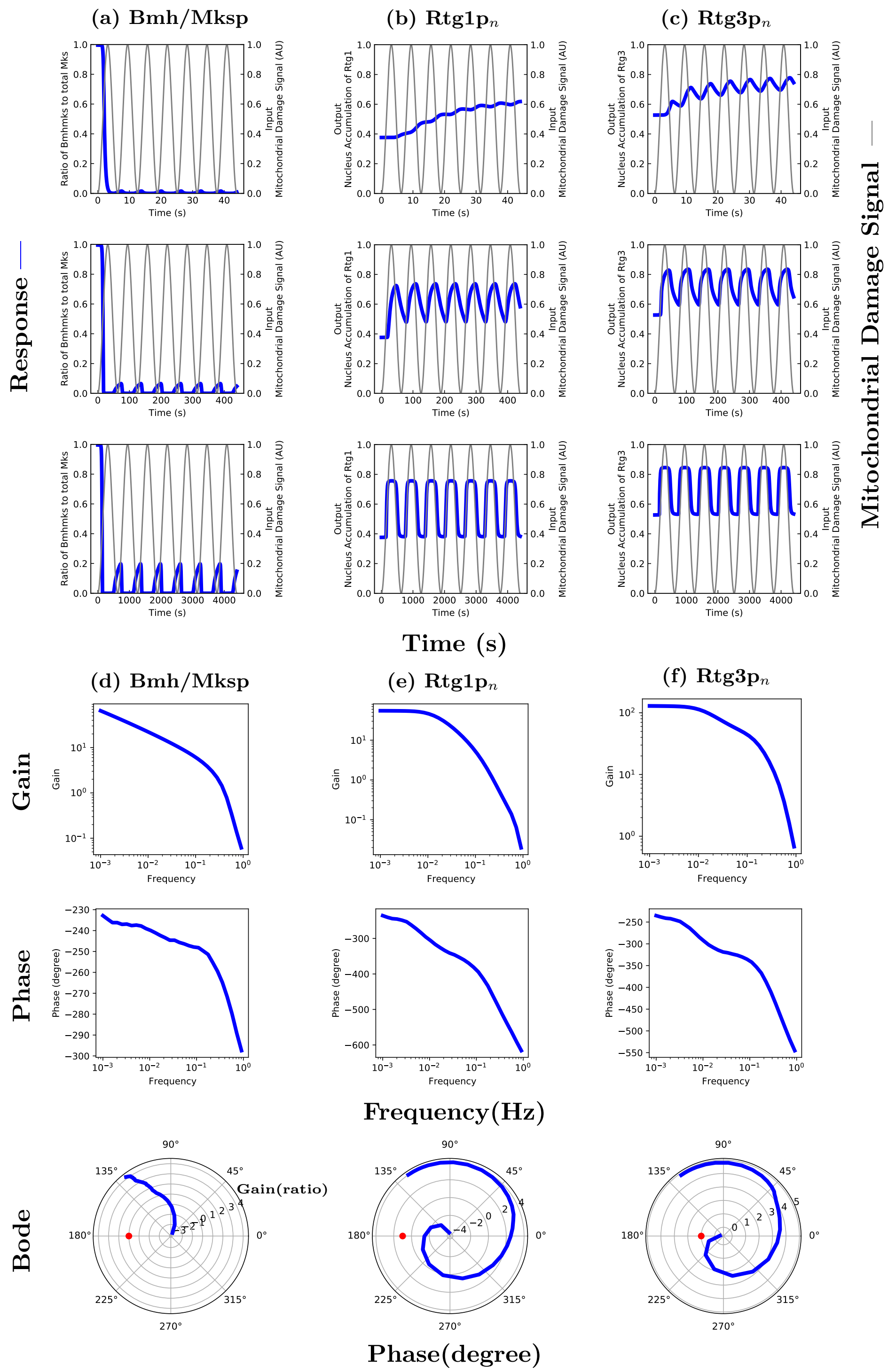

**Fig S4. Frequency analysis of Bmhp/Mksp, Rtg1p, and Rtg3p with respect of mitochondrial damage.**
(a-c)The frequency response of mitochondrial damage signal and RTG protein concentrations. The sinusoidal inputs (thin gray curves) with $1.6 \times 10^{-3}$ Hz, $1.6 \times 10^{-2}$ Hz and $1.6 \times 10^{-1}$ Hz reveals the RTG signaling pathway as a low-pass filter. The y-axis represents nuclear accumulation induced by the input signal, and the quantity is described by the ratio of nuclear concentration to total protein concentration. (d-f) Nyquist plots. In the Nyquist plots, red dot represents the critical point (1 + 0j), and the responses are represented as blue dots. The gain is measured when the oscillation reaches to stationary phase.

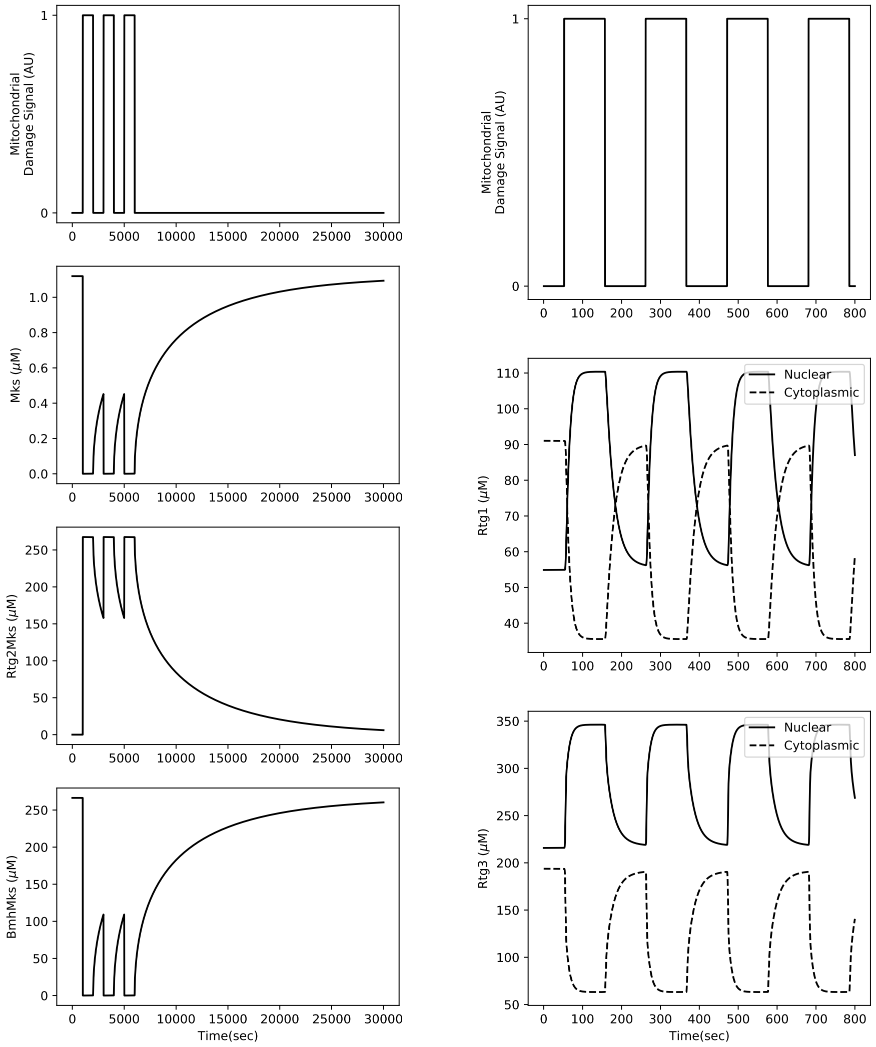

**(a)** **(b)**

**Fig S5. The square response of the proposed RTG model.**
(a) The square response of Mksp with its heterodimers, Bmh/Mksp and Rtg2mksp, reveals the sign-sensitive delay pattern, where the delay depends on the sign of the ON or OFF step. In Mksp, the sign-sensitive delay can protect against input fluctuations. The high binding coefficient of Mksp and its partners, Bmhp and Rtg2p, results in low concentrations of unbound Mksp. (b) Steady-state abundance of Rtg1/3p in the nucleus and cytosol in response to the mitochondrial damage. The nuclear accumulation of Rtg1p and Rtg3p induced by mitochondrial damage with square wave signal. Unlike Mksp, The square response delay of Rtg1/Rtg3p is insensitive to the sign of the step. The Rtg1p and Rtg3p accumulate in the nucleus when the damage signal turns ON. Note, the nuclear concentration of Rtg3p is always higher than the cytoplasmic concentration regardless of whether the mitochondrial damage signal is ON or OFF [2].

| Table of RTG-associated Proteins | | | |
|---|---|---|---|
| Standard Name | Systematic Codes | Function | Human homolog* |
| Rtg1 / Rtg3 | YOL067C / YBL103C | Activates retrograde response | MYC-MAX |
| Rtg2 | YGL252C | Promotes translocation of Rtg1-Rtg3 from cytosol to nucleus; binds with Mks1 | unclear |
| Mks1 | YNL076W | Phospholates Rtg3; downregulates RTG pathway | MKS1 |
| Bmh1 | YER177W | Binds with Mks1, and forms Bmh1/2-Mks1 to phosphate Rtg3 | YWHAE |

**Table S1. Proteins associated with the yeast mitochondrial retrograde response.** *Human homologs are also referenced. [1, 9, 18]

| Mitochondrial Damage Signal | Existence of Rtg3p | Existence of Rtg2p | Nucleus Accumulation of Rtg1p [1] |
|---|---|---|---|
| $> 0$ | $> 0$ | $= 0$ | No |
| $> 0$ | $= 0$ | $= 0$ | No |
| $> 0$ | $= 0$ | $> 0$ | No |
| $= 0$ | $> 0$ | $> 0$ | Yes |
| $= 0$ | $> 0$ | $= 0$ | Yes |
| $= 0$ | $> 0$ | $> 0$ | Yes |
| $= 0$ | $= 0$ | $= 0$ | Yes |
| $= 0$ | $= 0$ | $> 0$ | Yes |

**(a)** Translocation of Rtg1-GFP

| Mitochondrial Damage Signal | Existence of Rtg1p | Existence of Rtg2p | Nucleus Accumulation of Rtg3p [1] |
|---|---|---|---|
| $> 0$ | $> 0$ | $= 0$ | No |
| $> 0$ | $= 0$ | $= 0$ | Yes |
| $> 0$ | $= 0$ | $> 0$ | Yes |
| $= 0$ | $> 0$ | $= 0$ | No |
| $= 0$ | $> 0$ | $> 0$ | No |
| $> 0$ | $> 0$ | $> 0$ | Yes |

**(b)** Translocation of Rtg3-GFP

| Mitochondrial Damage Signal | Existence of Mksp | Existence of Rtg2p | Nucleus Accumulation of Rtg1p and Rtg3p [6] |
|---|---|---|---|
| $> 0$ | $= 0$ | $= 0$ | Yes |
| $> 0$ | $= 0$ | $> 0$ | Yes |

**(c)** Translocation of Rtg1-GFP and Rtg3-GFP in $\Delta$*mks* strain.

**Table S2. Properties of Rtg1/3p nucleus translocation.**
0, deleted proteins; $> 0$, wild type proteins.

| Kinetic Coeff. | Description | Value (AU) |
|---|---|---|
| $n^s$ | Hill coefficient of input signal | 7 |
| $k_{sV}$ | Maximum activation coefficient of Rtg2p | $1.17 \times 10^1$ |
| $k_{sd}$ | Dissociation constant of Rtg2p activation | $9.65 \times 10^{-1}$ |
| $k_{ri}$ | Inactivation coefficient of Rtg2p | 4.95 |
| $k_{rM}$ | Binding coefficient of active Rtg2p and Mks | $1.60 \times 10^3$ |
| $kn_{rM_i}$ | Degradation coefficient of Rtg2/Mksp heterodimer | $4.25 \times 10^{-2}$ |
| $k_{bm}$ | Binding coefficient of Bmhp and Mksp | $5.90 \times 10^{-2}$ |
| $k_{bmi}$ | Degradation coefficient of Bmh/Mksp heterodimer | 2.41 |
| $k_I^{13}$ | Maximum inactivation rate on Rtg1/Rtg3p heterodimer | $9.23 \times 10^{-2}$ |
| $k_{V_I}^{13}$ | Maximum rate of Rtg3p inactivation | $2.48 \times 10^3$ |
| $k_{d_I}^{13}$ | Dissociation constant of Rtg3p inactivation | $2.51 \times 10^1$ |
| $k_{c_A}^3$ | Kinetic coefficient of Rtg3p activation in cytosol | $3.07 \times 10^1$ |
| $k_{c_I}^3$ | Kinetic coefficient of Rtg3p inactication in cytosol | $1.25 \times 10^{-2}$ |
| $k_{n_I}^3$ | Kinetic coefficient of Rtg3p inactivation in nucleus | 1.49 |
| $k_c^{13}$ | Kinetic coefficient of the binding of Rtg1p and Rtg3p in cytosol | $2.34 \times 10^2$ |
| $k_{c_n}^{13}$ | Kinetic coefficient of Rtg1/3p degradation in cytosol | $2.21 \times 10^2$ |
| $k_n^{13}$ | Kinetic coefficient of the binding of Rtg1p and Rtg3p in nucleus | $4.61 \times 10^2$ |
| $k_{n_n}^{13}$ | Kinetic coefficient of Rtg1/3p degradation in nucleus | $5.96 \times 10^{-1}$ |
| $k_{in}^1$ | Kinetic coefficient of Rtg1p nucleus translocation from cytosol | 3.20 |
| $k_{out}^1$ | Kinetic coefficient of Rtg1p cytoplasmic translocation from nucleus | $5.97 \times 10^3$ |
| $k_{inA}^3$ | Kinetic coefficient of the nucleus translocation of active Rtg3p | 1.02 |
| $k_{outA}^3$ | Kinetic coefficient of the cytoplasmic translocation of active Rtg3p | $4.90 \times 10^{-2}$ |

| | | |
|---|---|---|
| $k^3_{inI}$ | Kinetic coefficient of the nucleus translocation of inactive Rtg3p | $3.80 \times 10^{-2}$ |
| $k^3_{outI}$ | Kinetic coefficient of the cytoplasmic translocation of inactive Rtg3p | $1.23 \times 10^{-1}$ |
| **Protein** | | **Concentration (AU)** |
| $Rtg2^{ina}_c$ | Inactivated Rtg2p in cytosol | $3.76 \times 10^2$ |
| $Rtg2^{act}_c$ | Activated Rtg3p in cytosol | 8.78 |
| $Mks$ | Mksp protein (system name: Mks1p) | $8.07 \times 10^{-4}$ |
| $Rtg2Mks_c$ | Heterodimer of Rtg2p and Mksp in cytosol | $2.67 \times 10^2$ |
| $Bmh$ | Bmhp protein (including Bmh1p and Bmh2p) | $10.00 \times 10^3$ |
| $BmhMks$ | Heterodimer of Bmhp and Mksp | $1.97 \times 10^{-1}$ |
| $Rtg13^{act}_c$ | Heterodimer of Rtg1p and activated Rtg3p in cytosol | $2.57 \times 10^1$ |
| $Rtg13^{ina}_c$ | Heterodimer of Rtg1p and inactivated Rtg3p in cytosol | $2.14 \times 10^1$ |
| $Rtg3^{ina}_c$ | Inactivated Rtg3p in cytosol | $1.71 \times 10^1$ |
| $Rtg3^{act}_c$ | Activated Rtg3p in cytosol | $2.49 \times 10^1$ |
| $Rtg3^{act}_n$ | Activated Rtg3p in cytosol | $1.66 \times 10^1$ |
| $Rtg3^{ina}_n$ | Inactivated Rtg3p in nucleus | $2.06 \times 10^2$ |
| $Rtg1_c$ | Rtg1p in cytosol | 1.06 |
| $Rtg1_n$ | Rtg1p in nucleus | $5.67 \times 10^{-4}$ |
| $Rtg13^{act}_n$ | The heterodimer of Rtg1p and activated Rtg3p in nucleus | 7.31 |
| $Rtg13^{ina}_n$ | The heterodimer of Rtg1p and inactivated Rtg3p in nucleus | $9.04 \times 10^1$ |

**Table S3. Parameter set of the proposed model.**
The table includes the kinetic coefficient and the steady-state concentrations of RTG proteins categorized by the activation state, location and the form of heterodimer. The parameter fulfills the conditions listed in Table S2, and is validated by Figure 2. in main text (A.U. represents arbitrary unit).

2. Liu Z, Sekito T, Epstein CB, Butow RA. RTG-dependent mitochondria to nucleus signaling is negatively regulated by the seven WD-repeat

protein Lst8p. EMBO Journal. 2002;20(24):7209–7219. doi:10.1093/emboj/20.24.7209.

3. Jazwinski SM. The retrograde response: When mitochondrial quality control is not enough. Biochimica et Biophysica Acta - Molecular Cell Research. 2013;1833(2):400–409. doi:10.1016/j.bbamcr.2012.02.010.
4. Knorre D, Sokolov S, Zyrina A, Severin F. How do yeast sense mitochondrial dysfunction? Microbial Cell. 2016;3(11):401–408. doi:10.15698/mic2016.11.537.
5. Zhang F, Pracheil T, Thornton J, Liu Z. Adenosine Triphosphate (ATP) Is a Candidate Signaling Molecule in the Mitochondria-to-Nucleus Retrograde Response Pathway. Genes. 2013;4(1):86–100. doi:10.3390/genes4010086.
6. Sekito T, Liu Z, Thornton J, Butow RA. RTG-dependent mitochondria-to-nucleus signaling is regulated by MKS1 and is linked to formation of yeast prion [URE3]. Molecular biology of the cell. 2002;13(3):795–804. doi:10.1091/mbc.01-09-0473.
7. Dilova I, Aronova S, Chen JCY, Powers T. Tor signaling and nutrient-based signals converge on Mks1p phosphorylation to regulate expression of Rtg1p Rtg3p-dependent target genes. Journal of Biological Chemistry. 2004;279(45):46527–46535. doi:10.1074/jbc.M409012200.
8. Ferreira JR, Spírek M, Liu Z, Butow RA. Interaction between Rtg2p and Mks1p in the regulation of the RTG pathway of Saccharomyces cerevisiae. Gene. 2005;354(1-2 SPEC. ISS.):2–8. doi:10.1016/j.gene.2005.03.048.
9. Jazwinski SM, Kriete A. The yeast retrograde response as a model of intracellular signaling of mitochondrial dysfunction. Frontiers in Physiology. 2012;3 MAY(May):1–12. doi:10.3389/fphys.2012.00139.
10. Butow RA, Avadhani NG. Mitochondrial signaling: The retrograde response. Molecular Cell. 2004;14(1):1–15. doi:10.1016/S1097-2765(04)00179-0.
11. Wang ZX. An exact mathematical expression for describing competitive binding of two different ligands to a protein molecule. FEBS Letters. 1995;360(2):111–114. doi:10.1016/0014-5793(95)00062-E.
12. Legewie S, Blüthgen N, Herzel H. Quantitative analysis of ultrasensitive responses. FEBS Journal. 2005;272(16):4071–4079. doi:10.1111/j.1742-4658.2005.04818.x.
13. Ferrell JE, Ha SH. Ultrasensitivity part I: Michaelian responses and zero-order ultrasensitivity; 2014.
14. Rackauckas C, Nie Q. DifferentialEquations.jl – A Performant and Feature-Rich Ecosystem for Solving Differential Equations in Julia. Journal of open research software. 2017;5. doi:10.5334/jors.151.

15. Ghaemmaghami S, Huh WK, Bower K, Howson RW, Belle A, Dephoure N, et al. Global analysis of protein expression in yeast. Nature. 2003;425(6959):737–741.

16. Gomar-Alba M, Morcillo-Parra MA, Olmo MLD. Response of yeast cells to high glucose involves molecular and physiological differences when compared to other osmostress conditions. FEMS Yeast Research. 2015;15(5):fov039. doi:10.1093/femsyr/fov039.

17. Delmans M, Hemberg M. Discrete distributional differential expression (D3E) - a tool for gene expression analysis of single-cell RNA-seq data. BMC Bioinformatics. 2016;17(1):110. doi:10.1186/s12859-016-0944-6.

18. Rios-anjos RM, Camandona VDL, Bleicher L, Ferreira-junior JR. Structural and functional mapping of Rtg2p determinants involved in retrograde signaling and aging of Saccharomyces cerevisiae. 2017; p. 1–27. doi:10.1371/journal.pone.0177090.
